# Towards the Interpretability of Deep Learning Models for Multi-modal Neuroimaging: Finding Structural Changes of the Ageing Brain

**DOI:** 10.1101/2021.06.25.449906

**Authors:** Simon M. Hofmann, Frauke Beyer, Sebastian Lapuschkin, Ole Goltermann, Markus Loeffler, Klaus-Robert Müller, Arno Villringer, Wojciech Samek, A. Veronica Witte

## Abstract

Brain-age (BA) estimates based on deep learning are increasingly used as neuroimaging biomarker for brain health; however, the underlying neural features have remained unclear. We combined ensembles of convolutional neural networks with Layer-wise Relevance Propagation (LRP) to detect which brain features contribute to BA. Trained on magnetic resonance imaging (MRI) data of a population-based study (n=2637, 18-82 years), our models estimated age accurately based on single and multiple modalities, regionally restricted and whole-brain images (mean absolute errors 3.37-3.86 years). We find that BA estimates capture aging at both small and large-scale changes, revealing gross enlargements of ventricles and subarachnoid spaces, as well as white matter lesions, and atrophies that appear throughout the brain. Divergence from expected aging reflected cardiovascular risk factors and accelerated aging was more pronounced in the frontal lobe. Applying LRP, our study demonstrates how superior deep learning models detect brain-aging in healthy and at-risk individuals throughout adulthood.

## 1. Introduction

With the advent of large-scale magnetic resonance imaging (MRI) studies (e.g., *UK Biobank, Sudlow et al. 2015; LIFE, Loeffler et al. 2015*), the estimation of brain age (BA), and its contrast to the chronological age of a person (diverging BA, DBA), have become an increasingly predictive imaging marker for brain health. Higher DBA relates to accelerated cognitive decline, pathologies such as Alzheimer Disease (AD), hypertension and type 2 diabetes, as well as other lifestyle-related cardiovascular risk factors *(Franke and Gaser 2019; Dadi et al. 2020)*. However, underlying alterations of neural structures reflecting the relationship between BA and such factors are not well known. BA has been linearly estimated on predefined neuroimaging outcomes (e.g., cortical thickness maps *Liem et al. 2017*). Yet, feature extraction and preprocessing could lead to overconfidence w.r.t., or to the dismissal of, neural properties that can be relevant to BA. In contrast, deep learning (DL) models, specifically convolutional neural networks (CNNs; *LeCun et al. 1989; Ji et al. 2013*) are trained on *raw* data and provide more precise BA estimates *(Cole et al. 2017; Cole and Franke 2017)*. Particularly on large MRI datasets CNNs converge to a minimal mean absolute error (MAE) of 2.14 years *(Peng et al. 2021;* see also *Jonsson et al. 2019; Feng et al. 2020; Kolbeinsson et al. 2020; Dinsdale et al. 2021; Levakov et al. 2020; Bashyam et al. 2020)*. Despite these advantages, their complex architectures restrict straightforward interpretations of which image features drive their estimates, known as the *black-box problem (Samek et al. 2019; Samek et al. 2021)*. Several methods have been proposed to open the *black-box (Samek et al. 2021)*, such as perturbation and gradient techniques *(Baehrens et al. 2010; Simonyan et al. 2014; Zeiler and Fergus 2014; Sundararajan et al. 2017; Zintgraf et al. 2017; Smilkov et al. 2017)* , which also have been applied for BA predictions *(Levakov et al. 2020)*. While many of these methods highlight input areas or intermediate feature maps that are relevant for the prediction, they do not indicate whether this information increases or decreases the predictor output. For the continuous case of BA estimates this means that neither the pace of aging processes (i.e., DBA), nor the state of their progression (BA) can be inferred from computed saliency maps.

Conversely, the Layer-wise Relevance Propagation algorithm (LRP) highlights relevant areas in the input (image) that both favor and dismiss corresponding output decisions *(Bach et al. 2015; Montavon et al. 2018; Lapuschkin et al. 2019)*. LRP has been successfully used with DL in MRI-based classification tasks *(Böhle et al. 2019; Eitel et al. 2019; Thomas et al. 2019)*. However, the biological alterations that underlie aging are continuous in nature, which raises more challenges for both the DL model, and, consequently, its interpretation.

Here, we therefore aimed to provide a novel, openly available analysis pipeline extrapolating from a proof-of-concept simulation study to the implementation of superior CNNs on multi-modal MRI with the explanation algorithm LRP. Specifically, we asked which neurostructural features drive individual predictions and whether BA truly captures biological aging processes. On a group level we explored, how DBA is modulated by cardiovascular risk factors, and how this relationship manifests in distinct neural features. Based on previous findings, we hypothesized that BA relies on grey matter atrophy which include (pre)frontal and mesiotemporal cortex and cerebellum, and that risk factors such as obesity, hypertension and type 2 diabetes correlate with higher DBA, reflected in augmented vascular pathologies such as higher white matter lesion load. Importantly, opening the black box of DL image analysis is expected to reveal novel features of MRI-based neuronal properties that contribute to BA estimates, and thus advance our knowledge of brain health in aging.

## 2. Materials and methods

### 2.1. Data acquisition

The LIFE Adult study *(Loeffler et al. 2015)*, a population-based cohort study, encompasses dense clinical screenings of more than 10,000 participants coming from the area of Leipzig, Germany. Among others, the screening included measures of height, weight, blood pressure, blood-based biomarkers, cognitive performance and questionnaire batteries on mental health, and lifestyle (for more details see: *Loeffler et al. 2015*).

#### 2.1.1. Study sample and exclusion criteria

Of the more than 10,000 subjects of the LIFE Adult study, 2637 participants underwent a 1-hour MRI recording session at baseline. Of those participants with MR-scans, 621 participants were excluded mainly due to pathologies, leaving 2016 subjects for further analysis (age range 18-82 years, mean_age_ = 57.32, median_age_ = 63.0; n_female_ = 946; see **Fig. 2** and **Fig. A3** in **Appendix D**). Partially overlapping exclusion criteria were previous strokes (n=54), excessive brain lesions rated by trained medical staff (n=114), including white matter (WM) lesions rated with a Fazekas *(Fazekas et al. 1987)* score of 3 (n=44), radiological diagnosis of brain tumor (n=22), diagnosis of multiple sclerosis (n=5), epilepsy (n=27), cancer treatment in the last 12 months (n=109), centrally active medication (n=275), cognitive impairments indicated by a MMSE score < 26 (n=80), and poor quality MRIs (failing a visually quality check, e.g., regarding motion artefacts, n=41).

#### 2.1.2. MRI data

MRI data was acquired in a 1-hour recording session using a 32-channel head coil in a 3T Siemens Verio scanner. Various MRI sequences were applied (see *Loeffler et al., 2015*). For this study, we trained models on three MRI sequences used in clinical settings: i) structural T1-weighted images were taken with an MP-RAGE sequence (1 mm isotropic voxels, 176 slices, TR=2300 ms, TE=2.98 ms, TI=900 ms, field of view 256 x 240 x 176 mm^3^, sagittal orientation) which is often used to quantify cerebrospinal fluid, white and gray matter among others. ii) Fluid-attenuated inversion recovery images (FLAIR) were acquired (1 mm isotropic voxels, 192 slices, TR=5000 ms, TE=395 ms, TI=1800 ms, field of view 250 x 250 x 192 mm^3^, sagittal orientation). FLAIR is highly sensitive towards lesions in the WM, which are known to accumulate with age *(Tang et al. 1997; Ge et al. 2002; Beck et al. 2021)*. Lastly, iii) susceptibility-weighted magnitude images (SWI) are used to detect iron-deposits in the basal-ganglia *(Pfefferbaum et al. 2009; Bekiesinska-Figatowska et al. 2013)*, which could be linked to neurodegeneration and cognitive decline *(Haller et al. 2010; Du et al. 2018; Thomas et al. 2020)*, and are used to discover brain hemorrhages. SWIs were recorded with a T2*-weighted pulse sequence (0.8 x 0.7 x 2.0 mm non-isotropic voxels, 64 slices, TR=28 ms, TE=20 ms, field of view 230 x 173 x 128 mm^3^, sagittal orientation).

### 2.2. MRI preprocessing

MRIs of the three sequences (T1, FLAIR, SWI) were saved in three processing stages: *raw*, *freesurfer volume* (*recon-all*, *FreeSurfer 5.3.0*; *Fischl 2012*), and *MNI stage* (MNI152; *Fonov et al. 2011*, 2mm; via *ANTs 2.2*, *Tustison et al. 2020*). In the *freesurfer volume stage*, FLAIR and SWI images were linear registered (linear interpolation; *ANTs 2.2*) to the corresponding space of the T1-weighted images (‘brain.finalsurf.mgz’), which were subject to various intensity normalization steps and a skull stripping procedure, which are all part of the preprocessing steps in *FreeSurfer* (for more details see the **Appendix A**). For memory and processing efficiency, all images in all stages were pruned, i.e., their background was maximally removed, while keeping the same volume shape in the respective stage and, for raw images, respective sequence across all participants. These minimally-sized volumes were constrained to have a 2-voxel margin around the full brain of the largest brain in the whole dataset in the respective stage and sequence. Moreover, the image data of each subject was compressed by clipping upper intensity values to 383 (255 + 50%), which affected an insignificant number of voxels (< 0.001%), and subsequently, by re-normalizing the data between 0-255 (i.e., into 2^8^ discrete intensity values per voxel). The re-normalized images were then processed as memory efficient arrays of single-byte, unsigned integers (here: *uint8* type *numpy 1.18.1* arrays; *Harris et al. 2020*).

### 2.3. Prediction model architecture (MRI data)

Ensembles have been shown to predict more accurately and reduce model biases *(Dietterich 2000)*, also in the domain of BA prediction *(Jonsson et al. 2019; Couvy-Duchesne et al. 2020; Dinsdale et al. 2021; Peng et al. 2021; Levakov et al. 2020)*. The individual predictions of the base models were used to train and evaluate a linear head model of the respective sub - ensemble, leading to a weighted prediction of the whole ensemble. Subsequently, an additional linear top-head model was trained to aggregate predictions over those sub-ensembles (see the following paragraphs, and **Fig. 1**).

**Fig. 1.**
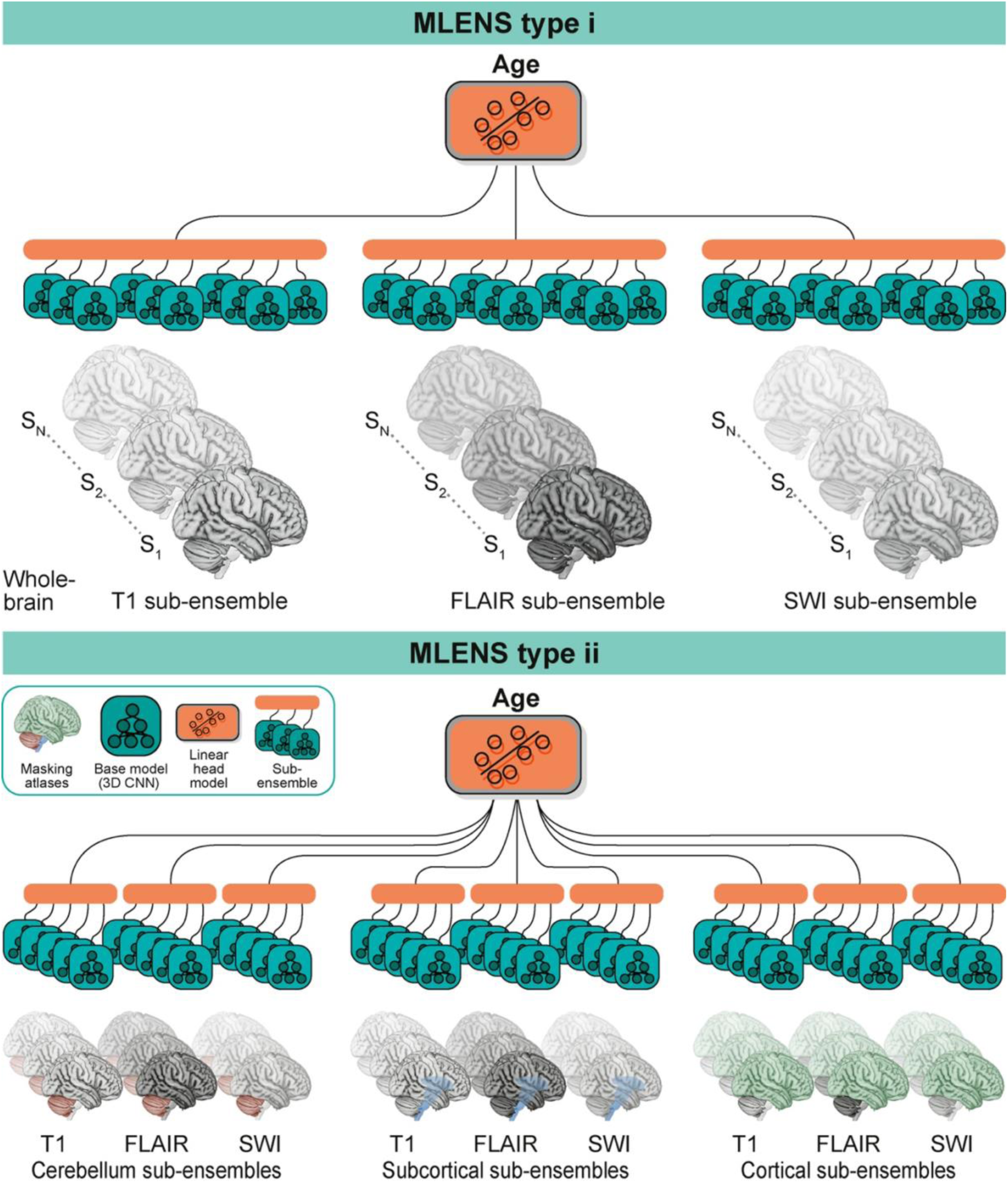
Multi-level ensembles (MLENS) MLENS trained on the different MRI sequences (T1, FLAIR, SWI; top: MLENS type i), and their combinations with 3 brain regions (bottom: MLENS type ii). The predictions of the sub-ensembles of each MLENS on the test set were used to train and evaluate the top-level linear head model.

#### 2.3.1. Base model

The base model architecture was a 3D convolutional neural network (3D-CNN; *LeCun et al. 1989; Lecun et al. 1998; Ji et al. 2013; Cole et al. 2017*), implemented in native *Keras 2.3.1 (Chollet 2015).* Base models were tested with two intermediate activation functions: i) the commonly applied rectified linear units (ReLUs), and ii) leaky ReLUs, which promise to overcome some of the drawbacks of absent gradients in standard ReLUs resulting from the background of MRIs, i.e., zero value input during training *(Maas et al. 2013)*. From bottom up, the network consists of 5 convolutional blocks (ConvB), each starting with a convolutional layer (n_filters_ , size_kernel_ ), followed by leaky ReLUs (, alpha = 0.2), and a 3D-max pool layer (, size_pool_ = 3^3^, stride = 2^3^). Then the signal was flattened to a 1-D vector, and during training a dropout layer (rate = 0.5) was applied. Finally, a fully connected layer (size = 64) with (leaky) ReLUs propagated the signal to the linear output neuron. The bias at the linear output layer was set to the target mean of the dataset (i.e., mean_age_ = 57.32), all other biases were randomly initialized around zero (*Keras’* default). The network was trained to minimize the mean squared error (MSE) w.r.t. chronological age, using the *ADAM* optimizer (learning rate = 5e^-4^; *Kingma and Ba 2015*). The data for the base models were split to a training, validation and test set (8:1:1 ratio). The training process on the training set was monitored on the validation set. The reported model performances are the results of its evaluation on the test set, and are given as the mean absolute error (MAE) for better interpretability.

#### 2.3.2. Model ensembles

Two types of multi-level ensembles (MLENS) were trained (**Fig. 1**): The first type consists of 3 sub-ensembles for 3 MRI sequences (T1, FLAIR, SWI), respectively. Each sub-ensemble has 10 base models (BM) that were independently trained on the same training data (whole brain data in *freesurfer volume stage* of its respective MRI sequence). Then, a linear head model (HM) with weight regularization, i.e. ridge regression (alpha = 1.) implemented in *scikit-learn 0.22.1 (Buitinck et al. 2013)*, was trained on the predictions *P* of the 10 BMs per sub-ensemble on the validation set (*P_val, BM_* = *X_train, sub-HM_* with shape: *N_val_* x 10; where *N_val_* is the number of samples in the validation set), and evaluated on the test set (shape of *X_test, sub-HM_* : *N_test_* x 10; where *X_test, sub-HM_* = *P_test, BM_*). The resulting predictions *P_test, sub-HMs_*of these 3 sub-ensembles on the test set were then used to train yet another head model on top of the MLENS in a 5-fold cross-validation (CV; *X_CV, top-HM_* = *P_test, sub-HMs_* of shape *N_test_* x 3) approach to obtain aggregated predictions across all MRI sequences. Note, only after the training of the whole MLENS, we evaluated single sub-ensembles, that is, we computed their MAE on the test set. This was done to compare the information gain between input MRI modalities with respect to age (see **Tab. 1**). Hence, using the test set predictions of the independently trained sub-ensembles for the training and evaluation of the MLENS top head model was still naïve about the corresponding participants’ age in the respective test fold of the CV, and only aimed to aggregate and weight the different input modalities.

**Table 1.**
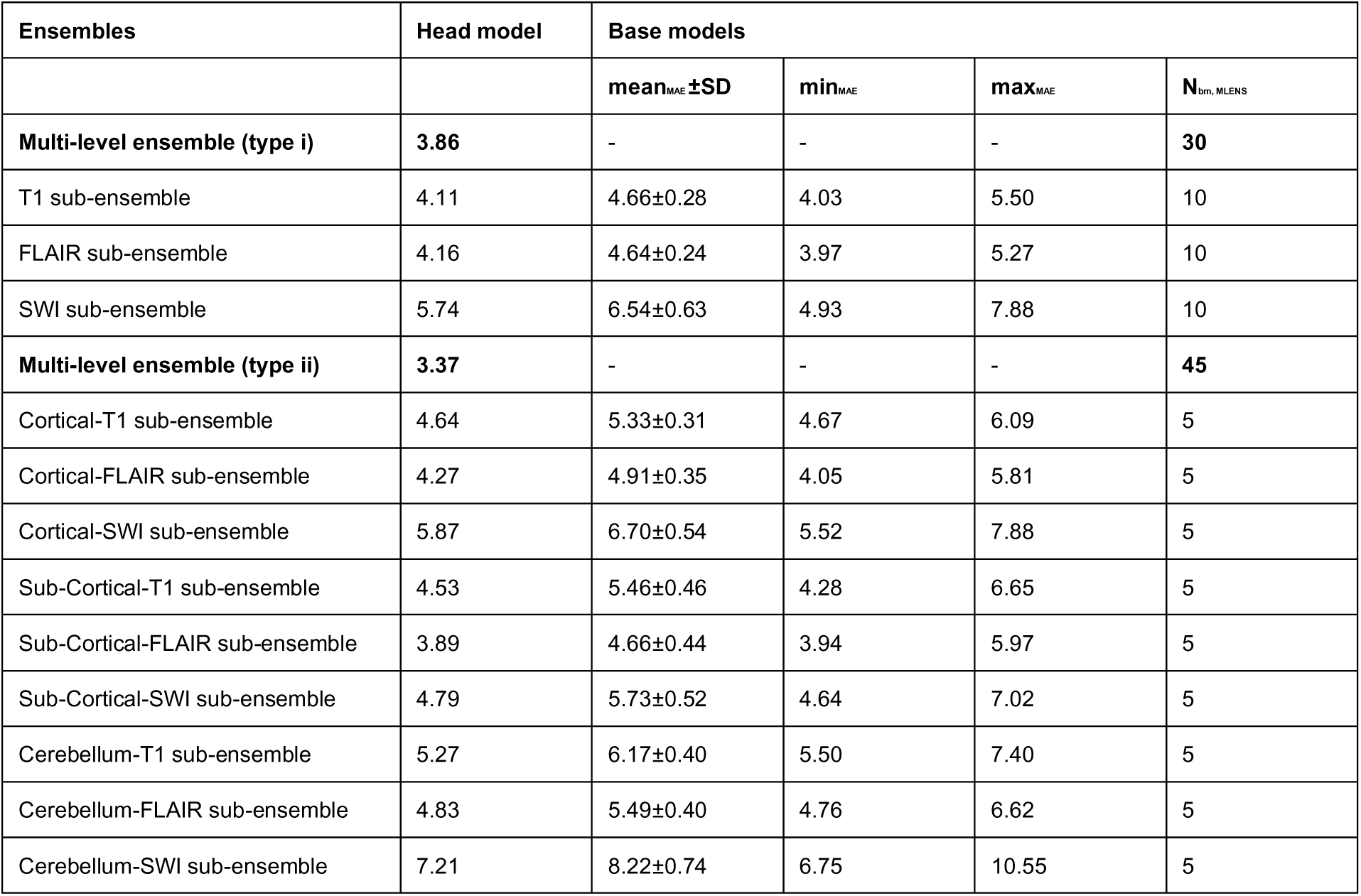
Prediction performances of both types of multi-level ensembles (MLENS type i, ii) and their respective sub-ensembles and 3D-CNN base models (bm), measured in mean absolute error (MAE). To receive an age estimate for each subject, MLENS were trained in a 10-fold cross-validation approach such that each subjects lies once in an unseen test set.

| Ensembles | Head model | Base models |  |  |  |
| --- | --- | --- | --- | --- | --- |
| | | mean<sub>MAE</sub> ± SD | min<sub>MAE</sub> | max<sub>MAE</sub> | N<sub>bm, MLENS</sub> |
| **Multi-level ensemble (type i)** | **3.86** | - | - | - | **30** |
| T1 sub-ensemble | 4.11 | 4.66±0.28 | 4.03 | 5.50 | 10 |
| FLAIR sub-ensemble | 4.16 | 4.64±0.24 | 3.97 | 5.27 | 10 |
| SWI sub-ensemble | 5.74 | 6.54±0.63 | 4.93 | 7.88 | 10 |
| **Multi-level ensemble (type ii)** | **3.37** | - | - | - | **45** |
| Cortical-T1 sub-ensemble | 4.64 | 5.33±0.31 | 4.67 | 6.09 | 5 |
| Cortical-FLAIR sub-ensemble | 4.27 | 4.91±0.35 | 4.05 | 5.81 | 5 |
| Cortical-SWI sub-ensemble | 5.87 | 6.70±0.54 | 5.52 | 7.88 | 5 |
| Sub-Cortical-T1 sub-ensemble | 4.53 | 5.46±0.46 | 4.28 | 6.65 | 5 |
| Sub-Cortical-FLAIR sub-ensemble | 3.89 | 4.66±0.44 | 3.94 | 5.97 | 5 |
| Sub-Cortical-SWI sub-ensemble | 4.79 | 5.73±0.52 | 4.64 | 7.02 | 5 |
| Cerebellum-T1 sub-ensemble | 5.27 | 6.17±0.40 | 5.50 | 7.40 | 5 |
| Cerebellum-FLAIR sub-ensemble | 4.83 | 5.49±0.40 | 4.76 | 6.62 | 5 |
| Cerebellum-SWI sub-ensemble | 7.21 | 8.22±0.74 | 6.75 | 10.55 | 5 |

For the second MLENS type, the MRI data (in *MNI stage, i.e.,* MNI152; see Section 2.2.) was additionally masked in three different brain regions defined by the three complementary atlases (see **Appendix A**: Brain atlases). For each combination of region and MRI sequence (3x3), 5 base models were trained, leading to a total of 45 base models. For each such combinatorial pair, its base model predictions were first aggregated with a linear head model (as above). Then, a linear top-head model combined these sub-ensemble predictions on the test set in the above mentioned 5-fold-cross-validation fashion to receive predictions across all input feature pairs (**Fig. 1**).

Both MLENS types (i, ii) can be conceptualized as *neural additive models (Hastie and Tibshirani 1990; Agarwal et al. 2020)*, i.e. sub-parts of the ensemble are trained on different input features.

To receive an age estimate for each subject, the training procedure was run 10 times, such that each subject lies once in the test set. In each of the runs the MLENS models were re-initialized.

### 2.4. Estimation of model uncertainty

Model certainty was measured subject-wise on both model levels, over each sub-ensemble and across them. That is, on the sub-ensemble level, model (un-)certainty is expressed as the standard deviation around the mean prediction of all its base models for each subject. Additionally, 95%-confidence intervals were computed for visual interpretation. Similarly, the standard deviation across the predictions of all sub-ensembles indicates the overall (un-)certainty of the MLENS. Note, the latter could also be interpreted as information gain across input features.

### 2.5. Prediction analyzer: Layer-wise Relevance Propagation

Layer-wise Relevance Propagation (LRP; *Bach et al. 2015; Montavon et al. 2018; Lapuschkin et al. 2019*) is an algorithm that provides explanatory heatmaps in the input to machine learning models, including non-linear deep learning models. To this end, the method decomposes the prediction *f*(*x*) of the model *f* w.r.t. the input *x* into relevance scores *R*. For deep learning models, this decomposition is computed layer-by-layer down to the input space, while satisfying the conservation criterion: ∑ Σ = *f*(*x*) <u>(for details, see</u> *Montavon et al. 2019*). In contrast to gradient-based and occlusion-based explanation methods, LRP is computationally efficient, since it only needs a single backward sweep. This is particularly important for large size MRI data. Moreover, LRP does not suffer problems such as *shattered gradients* or *pathological minima (Montavon et al. 2019; Samek et al. 2021)*, while it shows similarities to the benefits of gradient smoothing as in *SmoothGrad (Smilkov et al. 2017)*. In contrast to standard occlusion-based explanation methods, LRP takes not only local but also global feature interactions into account that are crucial for the model prediction (however, there can be occlusion-based methods formulated that overcome this locality issue; see *Samek et al. 2021)*. Lastly, Arras et al. (2022) could show in a ground-truth testbed developed for explanation methods that LRP performed best across 10 different algorithms.

LRP, which has been mainly employed in classification tasks, can be simply adapted to a regression problem *(Letzgus et al. 2022)*. Artificial neural networks (ANNs) used for classification usually have an output neuron for each class label in the employed dataset. LRP allows tracing the activation of each of these individual output neurons back to the input space through the network layers following a set of rules that depend on the layer types (for details, see *Montavon et al. 2019*). That is, LRP highlights areas in the input most relevant for the activation of the output neuron of interest (e.g., the neuron representing the ground-truth label, or the neuron with the highest output representing the model prediction). Information for the prediction will result in positive relevance scores in the input, while negative scores reflect information that the model considers as speaking against the respective output label. This feature of discretizing between positive and negative evidence makes LRP an useful approach among other explanation algorithms (e.g., *SmoothGrad*; *Smilkov et al. 2017*) of which many are used in absolute terms, i.e., without discretizing between positive and negative relevance (e.g., in *Levakov et al. 2020*).

ANNs for regression problems, mostly have only one output neuron (or more in multivariate regressions). In our case, adopting LRP for brain-age predictions means applying its algorithm starting at the single output neuron of the regression model. This is analogous, and mathematically equivalent to choosing the output neuron representing the ground-truth label of a given sample in a classification task. Differences are the task-specific objective function, the bias at the output layer, which we set to the distribution mean, and the accompanying interpretation of the relevance maps. Setting the output bias to the sample mean entails that positive relevance values indicate information towards the upper bound of the regression domain, while negative relevance values indicate the opposite (here: model evidence for a younger age).

#### 2.5.1. Simulation study on LRP for regression

We created two-dimensional images of tori on black backgrounds at an intensity range comparable to T1-weighted MRIs that exhibited inner and outer surface atrophies as a linear function of their *age (20-80 years)* with a normally distributed variance, to simulate cortical atrophy and enlargement of cerebrospinal fluid space. Additionally, we simulated that the *older* a torus was, the more *lesions* it accumulated within its body, appearing as image hyperintensities. In contrast to the atrophies, this accumulation of lesions was non-linearly increasing with age (i.e., onset of linear increase at age 40), also with a normally distributed variance. For each torus, the location of atrophies and lesions were known allowing for the evaluation of the sensitivity of the model represented in the relevance maps (see Section 2.5.2.). For the image details, please see the openly available code (https://github.com/SHEscher/XDLreg). We created 2000 tori, with a similar *age*-distribution as in the LIFE MRI sample. On this dataset, we then trained a 2D-version of the CNN as described above. Finally, LRP heatmaps were created on samples of the corresponding test set similar as described in following section. Since these heatmaps served only a qualitive analysis, we did not run statistical tests between them as we did for the MRI case.

#### 2.5.2. LRP for the MRI-based multi-level ensembles

LRP was applied on the trained base models in one ensemble of each type (via *iNNvestigate 1.08*; *Alber et al. 2019*), using the best-practice, composite rules *(Montavon et al. 2019; Kohlbrenner et al. 2020)* of LRP for CNNs (alpha = 1) implemented in *iNNvestigate* as "*LRPSequentialPresetA*". Note that we ran the LRP analysis only on models with ReLU activation functions, as it is recommended in *iNNvestigate*.

For the evaluation of the heatmaps, we took the average of the various relevance maps across base models similar to *Levakov et al. (2020)*. For between-subject analyses, we warped the subject respective heatmaps to MNI space. Relevance map aggregations within each subject were performed subsequently. The contribution of individual brain-regions to the model prediction was evaluated by mapping the LRP heatmaps to the merged brain-atlas, and the Juelich histological atlas (see **Appendix A**: Brain atlases). Additionally, we ran significance tests on the relevance maps with *FSL 5.0.8* (*randomise* function; using 5000 permutations and threshold-free cluster enhancement, TFCE) to determine brain areas which were statistically relevant for the BA prediction *(Jenkinson et al. 2012)*. This was done, across all participants on their absolute aggregated relevance maps (one-sample t-test). Absolute relevance values were taken, since they reflect meaningful information for a model, irrespective of the age of a participant’s brain; conversely, relevance values of zero reflect areas in the image that the model *ignored* for its age estimates. Contrastive relevance maps (unpaired two-sample t-test) were computed in a young (age ≤ 40 years) versus elderly (age ≥ 60 years) group on their signed aggregated relevance maps. In older adults (age ≥ 50 years), we analyzed in which brain regions relevance is attributed as function of the diverging (or delta) BA (DBA := y_predicted-age_ - y_true-age_) independent of chronological age. That is, we ran a generalized linear model (GLM; *FSL 5.0.8*, *randomize)*, with relevance maps as regressand, and DBA as regressor, while controlling for age as covariate. Additionally, we explored the role of a pathobiological biomarker (see the following section for more details), specifically type 2 diabetes mellitus on the BA estimates within a wider, older age range (50-75 years), contrasting diabetics to healthy controls (unpaired two-sample t-test on their signed aggregated relevance maps). Lastly, to test whether the MLENS capture individual WM lesions, we followed a two-step approach. First, we calculated for each individual a WM lesion probability map using the Lesion segmentation toolbox *(Schmidt et al. 2012)*, and applied a threshold of 0.8. In a second step, we aligned these binarized WM lesion maps to our relevance maps. For participants with more than 30 WM lesion voxels, we calculated the average relevance per WM lesion voxel. If the MLENS were able to capture individual WM lesions and use them as an information source to predict higher age, the calculated average relevance for these voxels should be positive. To increase the sample size for all tests, we combined relevance maps from the validation and test set.

### 2.6. Brain-age as a biomarker

As an exploratory analysis, we correlated (Pearson’s *R*; *scipy* 1.4.1; *Virtanen et al. 2020*; Bonferroni-corrected) DBA with a set of variables known to relate to accelerated brain aging.

These included cardiometabolic risk factors (BMI, waist-to-hip-ratio, hyperlipidemia, hypertension, systolic blood pressure, type 2 diabetes, glycated hemoglobin), genetic factors (apolipoprotein epsilon 4 risk-allele, *APoE4*, which has been associated with AD; *Strittmatter et al. 1993*), gender, time of education, cognitive functioning (composite score of executive functions, memory and processing speed, as reported in *(Kharabian Masouleh et al. 2016; Zhang et al. 2018)*, and neural integrity (here measured as the logarithm of the ratio between number of lesions and white matter volume). For this, we applied an overlapping sliding window approach over the full age range (width 10 years) to model age-related associations between DBA and the above-mentioned variables, and to minimize the effect of age on the prediction error itself. In each window we calculated *R* between DBA and the respective continuous variable. For the remaining categorical variables, which are all binary, Pearsons *R* is equivalent to the *Phi* coefficient or Kendall’s *Tau* coefficient that are usually applied for categorical variables. For simplicity, we report for both binary and continuous variables the corresponding coefficient as *R*. To control for multiple comparisons, we applied Bonferroni correction taking the number of variables into account (n=12).

## 3. Results

We implemented two types of **m**ulti-**l**evel **ens**embles (MLENS, **Fig. 1**) on three clinically relevant MRI modalities (T1-weighted, fluid-attenuated inversion recovery, FLAIR, and Susceptibility Weighted Imaging, SWI) of a well-characterized population-based cohort study (LIFE-Adult; *Loeffler et al. 2015*; age range 18-82 years, n = 2016).

Briefly, MLENS type I was trained on whole brain MRI with a sub-ensemble for each sequence with ten 3D-CNN models (base models, BM). Sub-ensembles served to extract information on model certainty and to compute more robust BA estimates. To additionally explore the contribution of three distinct brain regions (cortical, sub-cortical structures, and cerebellum) to the BA estimate, MLENS type ii was trained on 3^2^ combinations of the MRI sequences and the brain regions, while employing 5 BMs for each combination.

### 3.1. Model prediction performances

The MLENS type i had a MAE of 3.86 and performed slightly better than all its sub-ensembles (MAE_T1_ = 4.11, MAE_FLAIR_ = 4.16, MAE_SWI_ = 5.74; **Table 1**). The MLENS type ii had a smaller MAE of 3.37 (see **Fig. 2** for prediction accuracy and model uncertainty) and was again superior to the performances of its sub-ensembles (**Table 1**). Between both MLENS, there were highly significant correlations between their predictions (R = 0.97, p < 0.001) and their prediction errors (R = 0.73, p < 0.001) on the test set. Note that these models were trained with leaky rectified linear units (ReLUs), while models trained with standard ReLUs performed worse (MLENS type i, MAE = 3.88; MLENS type ii, MAE = 3.69; see **Appendix C Table A2**).

**Fig. 2.**
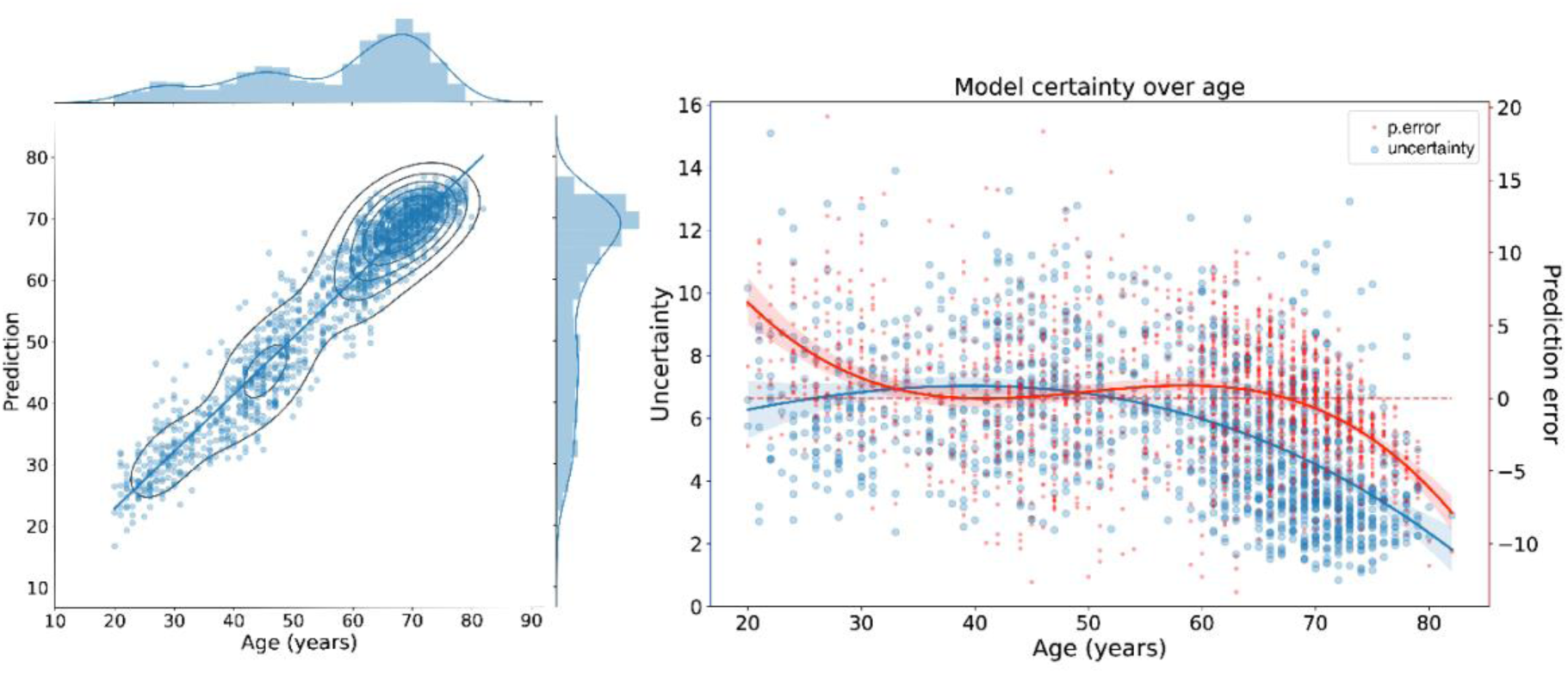
Brain-age prediction performance and model certainty of deep learning-based multi-level ensembles (MLENS) combining clinically relevant MRI sequences. Left panel: test set predictions of the MLENS type ii), trained on 3^2^ combinations of MRI sequences (T1, FLAIR, SWI) and brain regions (cortical, subcortical structures, and cerebellum). Right: prediction error (in red) and model uncertainty (in blue) per participant. Model uncertainty is measured as the standard deviation across the predictions of the sub-ensembles, and visualized as the width of the corresponding 95%-confidence interval. The modulation of both variables as function of age was modeled with a 3^rd^ order polynomial (red and blue lines). Both plots are produced over the concatenated test sets of the 5-fold-cross-validation, which were used to evaluate the top-level head models of the ensemble.

### 3.2. Relevance maps of model predictions

To verify the behavior of the LRP algorithm and its correct interpretation in a regression task, we first performed a simulation study.

The CNN model for the simulation task corresponded to a 2D-version of one base model in a MLENS. It was trained on a simulation dataset of abstracted head models (tori; **Fig. 3**), in which aging was simulated as the accumulation of *atrophies* and *lesions*. The model had a MAE of 2.80 on the hold-out test set. The prediction model captured the simulated aging process in both its facets well, which is revealed by the LRP relevance maps (i.e., heatmaps) highlighting the inner and outer borders (*atrophies*), and the added *lesions* within the *older* tori (30+ years; **Fig. 3**). Areas, where atrophies can occur were considered as information bearing, i.e., they received both positive and negative relevance. Moreover, the model seemed to cluster information w.r.t. its regression task, which is represented in the unique sign of relevance over larger areas (see both tori on the right, **Fig. 3**). That is, while there were accumulations of atrophies at the border of some tori, the CNN also took adjacent lesions into account to aggregate the overall information in a specific region. Note that in some occasions this could lead to inversely weighted relevance in single pixels or small areas (see upper left part of *green box* in **Fig. 3**). The sum over all distributed relevance *r* is a proxy for the final model prediction (). If it is positive, the prediction *p* is greater than the initiated model target bias (*b_t_*; here, set to the mean age of the sample: *b_t_* = 51.1 years), and vice versa for the negative case. Hence, the summed relevance represented the evidence over the whole image that the model accumulated to make its prediction.

**Fig. 3.**
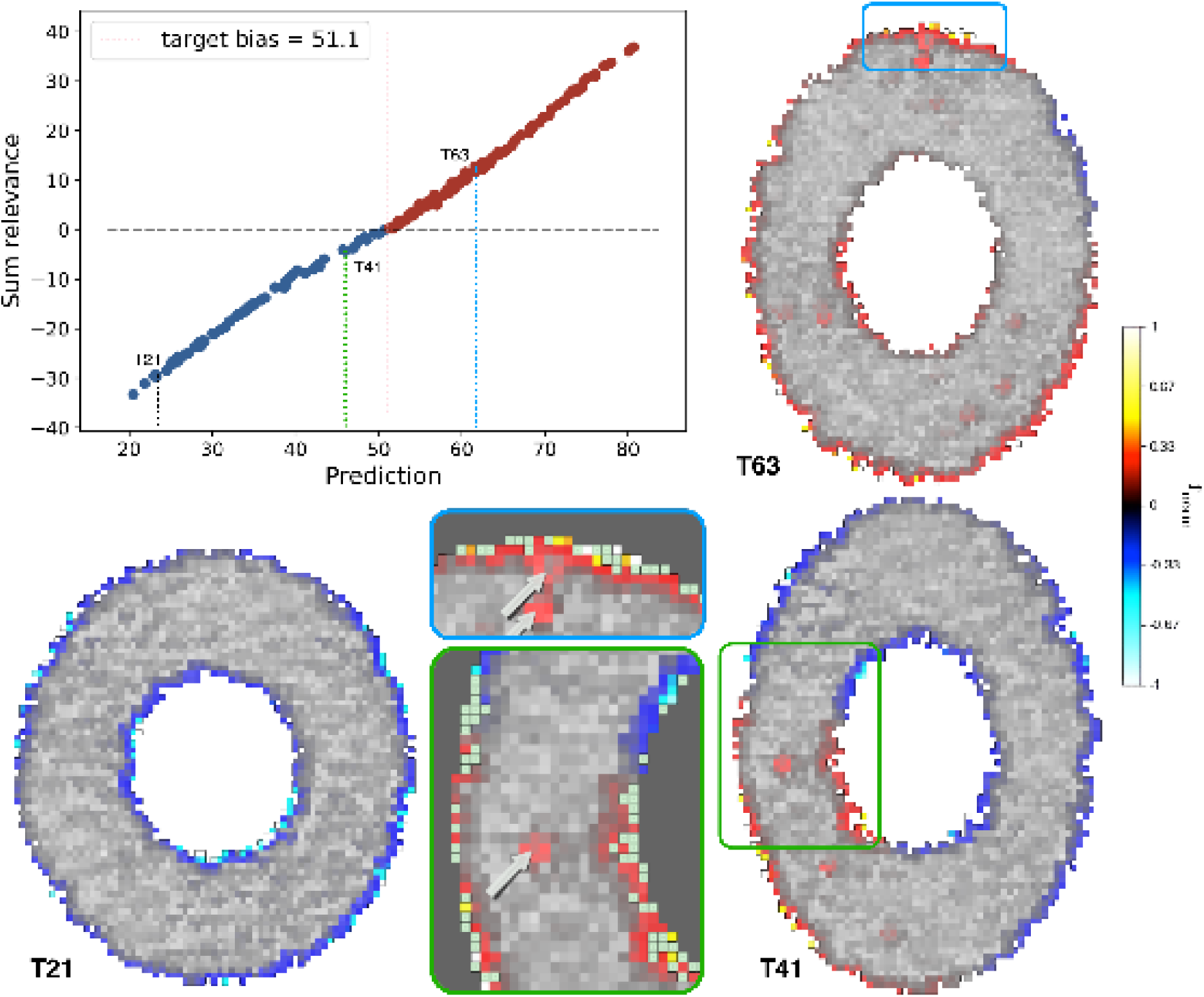
Analysis of simulated aging in artificial tori. *Top left*: Summed relevance per predicted sample in the test dataset, reflecting the model prediction relative to the sample mean (i.e., target bias = 51.1 years). That is, this global conversation property entails that all distributed relevance R in the input space of a given image x reflects the model prediction: , where R_i_ is the relevance at pixel i (Montavon et al., 2019). *Bottom left and right column*: three image samples of tori (T[age]) with their corresponding LRP relevance maps overlaid. *Gray boxes*: Details of relevance maps of tori T41 (green) and T63 (blue), respectively. Here, arrows indicate added lesions, while mint-green pixels at the inner and outer borders of the tori indicate ground-truth atrophies. Note that intact matter is predominantly attributed with negative (blue-turquois) relevance, indicating a younger age, while lesioned or atrophied matter is attributed with positive (red-yellow) relevance pointing to an older age. *Color coding*: relevance values were symmetrically clipped around zero at the 0.99-percentile, then normalized (r_norm_ ) and the corresponding colormap was multiplied by a factor of 5 for better contrasts. Note, while the model predictions are continuous, we deliberately decided for a binary color scaling to better contrasts the lower (young) and upper (old) bound of the regression.

#### 3.2.1. Relevance maps of the aging brain in individuals

Qualitative LRP analysis revealed individual relevance maps highlighting brain areas that *voted* for higher or lower BA predictions. Overall, we detected strong contributions from voxels in and around the ventricles and at the border from the brain to meningeal areas, independent of MRI sequence, while white-matter (WM) areas appeared to be less informative, except WM lesions in FLAIR images (**Fig. 4a**). In older participants, voxels covering cortical sulcal structures were often more relevant than in younger participants and *voted* more often in favor of older BA. Also, the corpus callosum, the brain stem and areas in and around the cerebellum appeared to be relevant structures, from which the models gained information for both younger and older participants. Overall, in all three major brain components (GM, WM, and cortical spinal fluids, CSF), there was a linear increase of relevance scores as function of age, being strongest in GM, and weakest in CSF (see **Fig. A4** and **Table A5** in **Appendix E**).

**Fig. 4.**
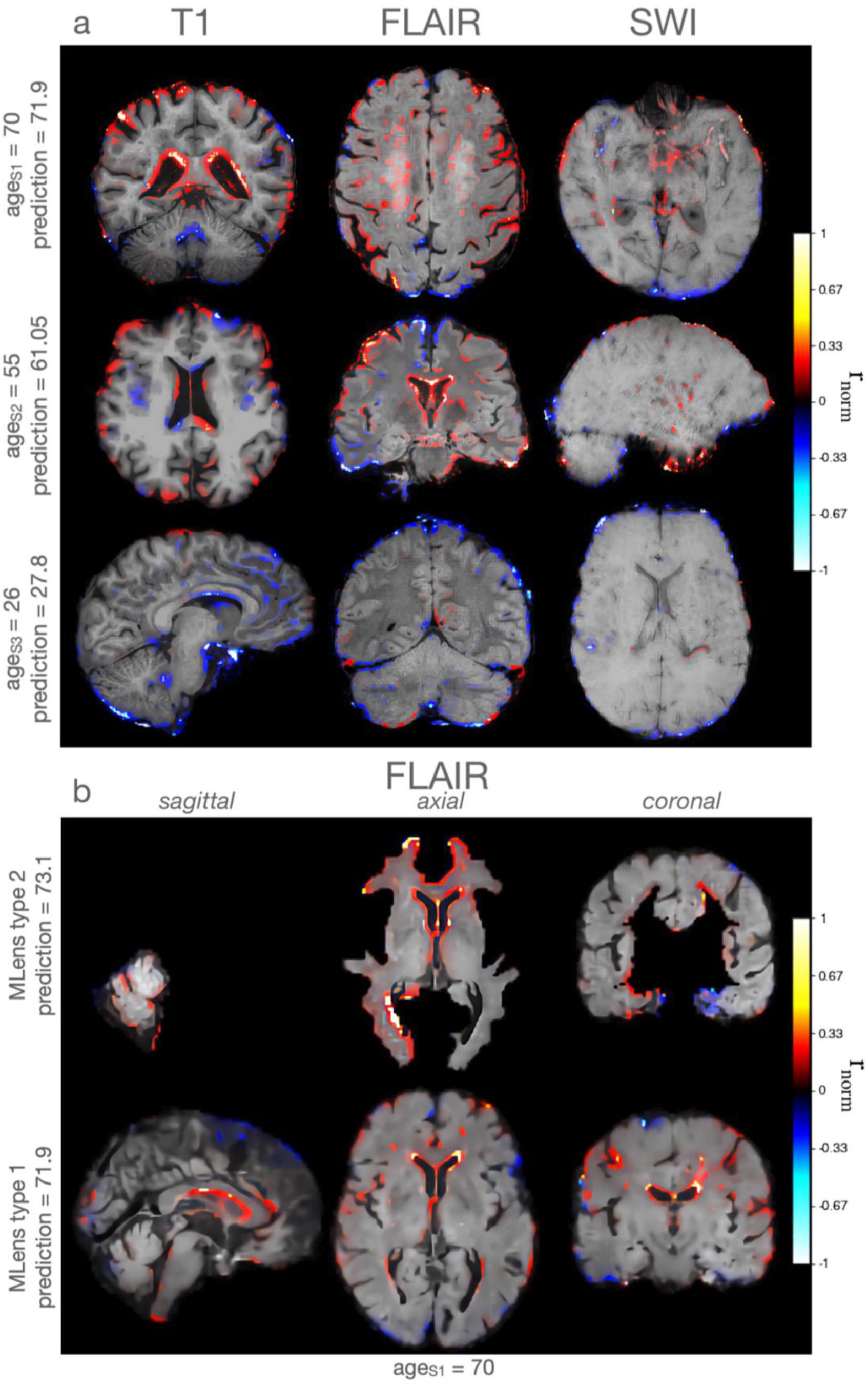
Exemplary individual LRP heatmaps. **(a)** of multi-level ensemble (MLENS) type i. trained on whole brain data. Rows: three participants (S1-S3) drawn from different age groups. Columns: three MRI sequences (T1, FLAIR, SWI), individually sliced in all three axes to highlight crucial areas that are unique to each participants’ age estimate. Next, to more global intact (mostly in young S3) or atrophied tissue (S1, S2), e.g., at the cortical surface, LRP also reveals smaller structures such as white-matter lesions (S1, FLAIR; which the model associates with higher age; see Section 3.2.2.), vessel expansions and putative small iron depositions, e.g., in the form of cerebral microbleeds (S1, SWI; see Discussion) driving the BA estimation. Relevance maps per subject were aggregated over the base models of each sub-ensemble. **(b) LRP heatmaps of regional (top row, type ii) and whole-brain (bottom row, type i) MLENS in elderly subject (S1)**. Here, models were trained on FLAIR data of cerebellum (left), subcortical structures (mid), and cortex (right), or of the whole-brain, respectively. For comparison, we warped the heatmap of whole-brain MLENS type i from subject space to MNI152 space (cf. top row in **a**). Note, that the average age of the cohort that was used to compute the MNI152 brain-space was 25.02±4.9 years (Fonov et al. 2011). Hence, the elderly subject S1 is warped to an aggregated young brain, which might lead to the impression that atrophies are less pronounced. Color coding: as in **Fig. 3**. Negative relevance scores (blue-turquois) represent model evidence in the input towards a younger age, and positive relevance (red-yellow) shows evidence towards a higher age.

Both types of MLENS (whole-brain type i and region-based type ii) found similar brain structures important for their prediction (**Fig. 4b**). Visually most recognizable are areas around the ventricles, and subject specific sulci, e.g., in the cortex and cerebellum.

#### 3.2.2. Statistical relevance maps over the adult lifespan

Quantitively, permutation-based one-sample t-tests (5000 permutations, threshold-free cluster enhancement, TFCE, and family wise error, FWE-corrected p ≤ 0.05) on combined relevance maps of the validation and test set (n_T1_ = n_FLAIR_ = 402, n_SWI_ = 314) of one MLENS type i revealed that on average, in all 3 MRI sequences, nearly the full brain contains meaningful information about BA (**Fig. 5**). The base models trained within the T1 sub-ensemble, gained most information in the lateral ventricle areas, corpus callosum, pre- and postcentral gyri in the motor and sensorimotor cortex, operculum, and all grey matter (GM) border areas including the frontal pole, temporal and visual poles and brainstem, and cerebellar borders. In the FLAIR sub-ensemble, most relevance was found around lateral ventricles, anterior temporal gyri, the pre- and postcentral gyri, and WM areas including cingulate gyrus, corpus callosum and fornix. Base models of the SWI sub-ensemble had a stronger focus on GM areas in the visual pole and occipital lobe, limbic areas, corpus callosum, WM fornix, internal capsule and on subcortical nuclei and brainstem areas, including striatum, subthalamic nucleus, raphe and substantia nigra. For an analysis of the differences between modalities see **Appendix F**.

**Fig. 5.**
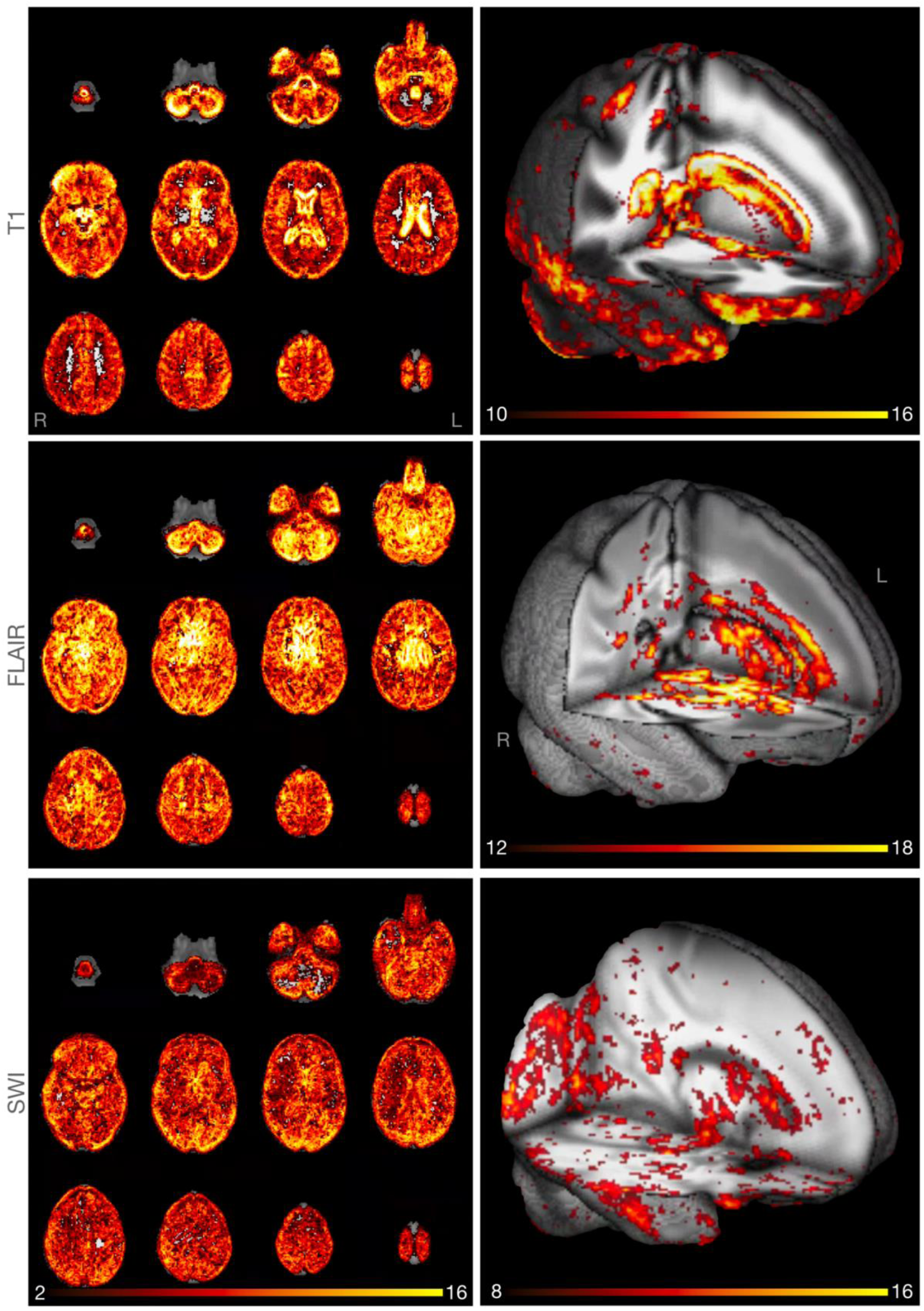
Relevant areas for brain-age predictions across adult lifespan. T-maps of one-sample t-test over aggregated, absolute relevance maps shown in several brain slices. *Left column*: t(2,16); MNI152 z-axis range: 3-74. The wider range of t-values shows that the model uses information from the whole brain for its age estimates. Right column: 3D-projection of t-maps focusing on higher t-values narrowly clipped for each MRI sequence, separately. These narrower t-maps highlight areas which dominate the model estimates. Top row: tested on the T1 sub-ensemble (Iype i; n = 402, t_max_ = 23.61). *Mid row*: FLAIR sub-ensemble (n = 402, t_max_ = 25.82). *Bottom row*: SWI sub-ensemble (n = 314, t_max_ = 16.07). The relevance scores were drawn from one of the MLENS type i models.

Next, we compared the LRP heatmaps of the young (age ≤ 40 years, n = 61) versus older cohort (age ≥ 60 years, n = 243). Areas showing greater relevance in older compared to younger brains (TFCE, FWE-corrected p ≤ 0.5) were found in the T1 sub-ensemble of MLENS type i in lateral ventricles, corpus callosum, amygdala, cerebral WM, particularly paracingulate gyrus, opercular cortex, and (secondary) somatosensory cortex. For FLAIR, there were increased relevance values found in cerebellum (specifically, left and right crus I-II), caudate, inferior frontal gyrus, pars triangularis, insular cortex, and inferior parietal lobule. For the SWI sub-ensemble, frontal pole, frontal orbital cortex, Inferior frontal gyrus, pars triangularis, precuneus, basal nuclei including putamen and caudate, and occipital pole showed higher (i.e., positive) relevance on average (**Fig. 6a**).

**Fig. 6.**
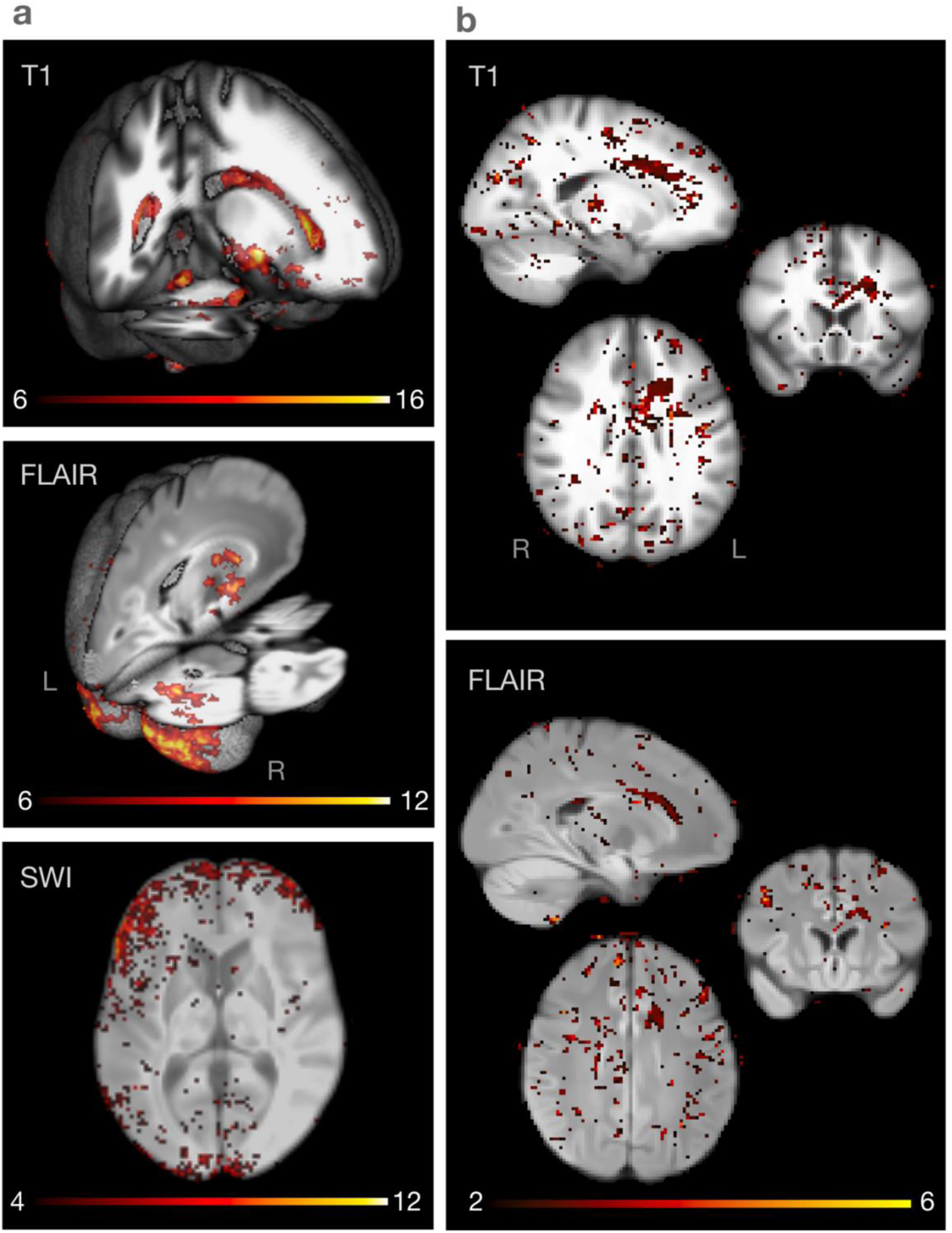
(a) Contrastive relevance maps of young vs. elderly participants. T-maps of two-sample t-test over relevance maps in the young and elderly group. Here, testing shows in which areas the relevance is greater in elderly (age ≥ 60 years) than in the young (age ≤ 40 years) group. Relevance maps were aggregated within each sub-ensemble of one of the MLENS type i models trained on T1 (top), FLAIR (mid), and SWI (bottom) data, respectively. **(b) Contrastive relevance maps of healthy vs. diabetic participants** T-maps (2, 6) of two-sample t-test show in which areas the relevance is greater in participants with type 2 diabetes than in healthy controls of the older cohort (60-75 years). Relevance maps were aggregated within each sub-ensemble of one of the MLENS type i models trained on T1 (top), and FLAIR (bottom). Note, only for T1 significant regional differences were found between the groups (see TFCE FWE-corrected maps in **Fig. A7** in the **Appendix G**). However, t-maps of T1 and FLAIR sub-ensembles show high correspondence (sliced in all three axes at x=-18., y=18.1, z=28.1).

#### 3.2.3. Lastly, based on our binarized WML probability maps, 654 participants had more than 30 WML voxels. The average relevance in WML voxels was significantly higher (i.e., 319 times) than the expected relevance per brain voxel (M_diff_ = 0.001, d = 0.9, t(653) = 22.95, p < 0.001).Relevance maps in diabetes and accelerated brain aging

To explore the role of health-related risk factors on BA, we contrasted the LRP relevance maps of subjects with type 2 diabetes (n=29) with healthy subjects (n=217) in the age range of 50 to 75 years (mean_age_ = 65.61). For the T1 sub-ensemble (MLENS, type i), clusters of higher positive relevance (non-healthy > healthy) were found to be significant in the pre- and postcentral gyrus near the cortico-spinal tract in the primary motor cortex (TFCE, FWE-corrected p ≤ 0.011), corpus callosum and cingulum (TFCE, FWE-corrected p ≤ 0.02; see **Fig. A7** in **Appendix G**). For the other two sub-ensembles (FLAIR, SWI), there were no clusters indicating significant regional differences. However, there was a high spatial correspondence between t-maps of the T1 and FLAIR sub-ensembles (**Fig. 6b**).

We further estimated the change in relevance maps as function of DBA, i.e., the signed prediction error, in an older cohort (age ≥ 50, mean_age_ = 67.07, n = 134), while controlling for age (as 2^nd^ order polynomial regression; cf. **Fig. 2)**. Accordingly, all clusters indicating a significant association spatially corresponded to areas found in the BA analysis, however, accelerated aging (DBA) was more strongly related to higher relevance values in specific regions (see **Fig. A8** in **Appendix G**): for the T1 sub-ensembles (MLENS type i) this difference was found in frontal pole, brain stem, outer cerebellar boarders, WM including the cortical spinal tract, putamen, caudate, amygdala, pre- and post-central gyri, and cingulate gyri. For the FLAIR sub-ensemble, primarily posterior region showed significant associations, including occipital and parietal pole, lingual gyrus, and cerebellum (crus I and II, V, VI). Finally, for the SWI sub-ensembles, posterior and anterior regions showed significant associations, including the frontal pole, frontal orbital cortex, occipital pole, cerebellum (crus I and II, vermis VIII), but also some more left-lateral parieto-temporal WM structures close to putamen and operculum (for all sub-ensembles; TFCE, FWE-corrected p ≤ 0.05).

### 3.3. Diverging brain-age and its relationship to other biomarkers

We found in the younger cohort (age < 45 years) that higher DBA correlated with cardiovascular risk factors such as hypertension and hyperlipidemia according to exploratory correlation analyses, which were run on the hold-out test sets (**Fig. 7**). In older subjects (age > 60 years) the most prevalent positive association of DBA was found with type 2 diabetes and accordingly, but weaker with glycated hemoglobin levels (HbA1c). BMI, waist-to-hip ratio and WM lesion load showed positive associations with DBA in participants almost across the full age range. Weak but relatively consistent trends appeared for the effect of gender (age > 30 years, where men had a higher BA on average) and the cognitive composite score, which showed a negative relationship with DBA. There was nearly no evident association between DBA and the presence or absence of an *Apolipoprotein E epsilon 4 gene allele* (APoE4), systolic blood pleasure or for higher education. Note, bivariate correlations were run in a sliding-age-window approach without adjusting for possible confounders; for multiple comparison a Bonferroni-correction was applied. While the here applied sliding-window approach aims to reduce the age-bias in DBA (**Fig. 2**), in a further analysis, we regressed this effect out over the full age range, before running the same correlation analysis (see **Appendix H**).

**Fig. 7.**
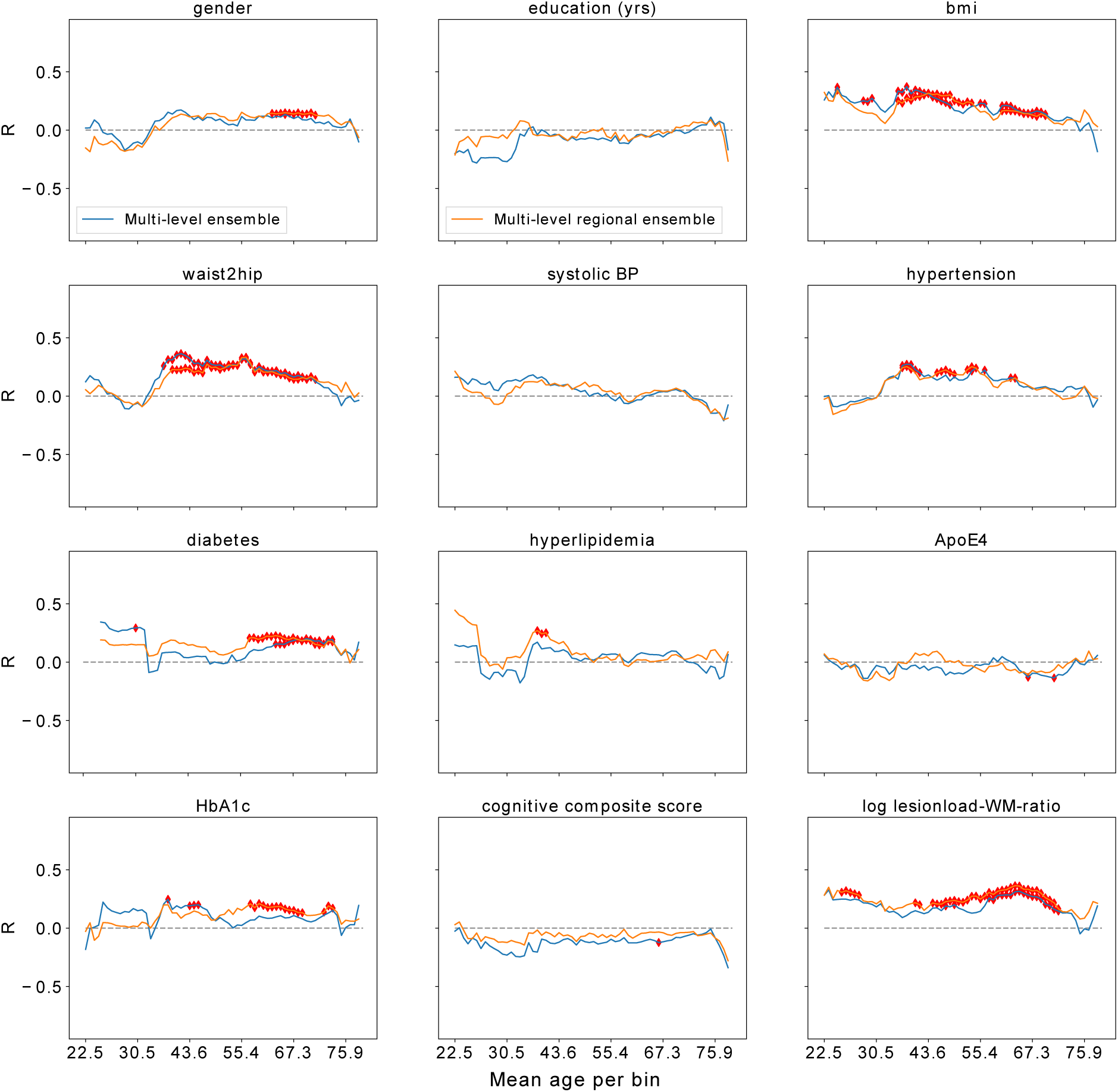
Relationship of diverging brain-age to biomarkers and lifestyle factors. Correlations (R) in overlapping sliding windows (width 10 years) between prediction errors (DBA) of both models (blue: type i; orange: type ii MLENS) and LIFE variables. Note, Kendall’s Tau for binary variables is equivalent to Pearson’s R that is used for the correlation analysis of continuous variables; we therefore name all correlations R for simplicity. Inverse width of the purple confidence band represents the number of participants per bin. *Red rhombus*: Bonferroni-corrected (n=12) p-value ≤ 0.05 per bin. Variables: education: time of education in years. bmi: body-mass-index. waist2hip: waist-to-hip-ratio. *systolic BP*: systolic blood pressure. *APoE4*: apolipoprotein epsilon 4 risk-allele carrier status. *HbA1c*: glycated hemoglobin. *Log lesionload-WM-ratio*: logarithmized ratio between number of lesions and white matter volume. Binary variables: *hypertension, diabetes, hyperlipidemia, APoE4*: no = 0, yes = 1. gender: female = 0, male = 1.

## 4. Discussion

The estimation of age and deviance from expected aging trajectories from brain images is a difficult task that has been solved to a surprisingly high accuracy using various DL architectures *(Cole et al. 2017; Cole and Franke 2017; Jonsson et al. 2019; Feng et al. 2020; Kolbeinsson et al. 2020; Dinsdale et al. 2021; Peng et al. 2021; Levakov et al. 2020; Bashyam et al. 2020).* We provide a further dimension to this challenge, namely, the means to extract insight from the trained neural networks, such that neurobiological theories can be validated and novel hypotheses can be generated. Specifically, we demonstrate that our accurate estimates of continuous brain aging can be related back to neurostructural features, by employing interpretable (here using LRP) DL-ensemble models on multi-modal 3D-MRIs that are trained end-to-end (i.e., no prior knowledge on brain features is induced to the model). Our analysis demonstrates that grey matter changes and atrophies detectable in the cortex, subcortex, cerebellum and brainstem, but also white matter lesions, as well as more global brain shrinkage represented in the larger size of ventricles and sulci drove the age estimates of the models. In further studies, this list of brain features should be validated further, for instance, by exploring the role of iron accumulations, cerebral microbleeds, or calcium depositions for the model estimates, which are associated with age and neural pathologies *(Haller et al. 2010; Du et al. 2018; Thomas et al. 2020)*. This happens to a degree that even parts of the brain and single MRI modalities (including SWI) led to accurate and comparable BA predictions. While voxels around the ventricular system and subarachnoid spaces were most informative for our model, the progression of aging and its pace (i.e., BA and DBA, respectively) could be related back nearly to the whole brain. Our simulation model furthermore revealed – as to be expected – that undamaged tissue (i.e., absence of atrophies and lesions) is associated with (young) age. From a methodological perspective, this demonstrated how the LRP algorithm can be integrated into a complex regression task on continuous aging, and how resulting relevance maps carry information about age-related changes. Moreover, we found that accelerated aging (DBA), which is associated with pathologies (here type 2 diabetes), shows relevant indicators in distinct brain areas, which could be differentiated by the complementary information from different MRI sequences and brain regions which we fed to the MLENS models, leading to overall better prediction results. With this, we established a novel DL-based pipeline for MRI analysis, which leverages the predictive advantages of this model class while at the same time making its estimates interpretable for research and clinical applications.

### 4.1. Opening the black-box of deep learning predictions

To understand the estimates of our DL models, we applied the LRP algorithm, which provides directed, i.e., sign-specific, relevance maps in the input space. Since, at the point of model inference a classification problem is mathematically similar to a regression problem, LRP could be straight-forwardly adapted to the purpose of our study (see Methods). We successfully validated this approach in the regression domain according to a simulation study with a 2D-version of the model architecture that we employed in the main study. We found that the DL model captures the simulated aging processes well by identifying the corresponding features. Explanations maps have to be interpretated carefully, avoiding potential confirmation biases of the researcher *(Adebayo et al. 2020)*. To validate the approach further, we ran additional simulations, where age is not modeled with strong image contrasts as in lesions and atrophies, but as function of shape and gradual local intensity shifts, respectively (for details see **Appendix B**). Also these analyses showed that the model captures the relevant information in the image, namely at the border of the torus for age estimations based on shape, and at the local area which was subject to age-related intensity shifts (**Fig. A1**). In the MRIs, we compared WM lesion maps with the relevance maps and found that also here, the model detects the WM lesions and associates higher age with them. LRP comes with the advantage of being directional, i.e., it indicates not only that a certain input area is relevant for a given prediction, but also whether it provides information in direction to the upper (here old age) or lower bound (young age) of the regression problem (**Fig. 3, 4**). The sign of the sum of relevance (SoR) is arbitrary in this case, essential is the magnitude of the value. Here, we chose to set the bias at the output layer of the CNNs to the mean of the target variable (age). As a consequence, the SoR becomes negative for predictions lower, and positive for estimates higher than the bias. The model does not only capture the features that represent the aging process (atrophies and lesions), but also the absence of them. That is, for the *young* torus it attributes (here negative) relevance also to its intact surface and borders. Moreover, LRP shows that the CNN finds irregular occurring features (here lesions) which were randomly placed. However, the interpretation of the local attribution of relevance needs to be taken with caution, as we observed that the model often generalizes relevance over larger areas of the simulated tori. One possible explanation for this is that relevance might be clustered over bigger areas after being passed through the intermediate pooling and convolutional layers in the network, which aggregate information over increasingly larger areas in the computed feature maps. Then, later layers (usually fully connected layers) make decisions over these pooled regions by attributing relevance towards one of the main directions in the regression *(Kohlbrenner et al. 2020)*.

### 4.2. Normal and accelerated brain aging

Applying LRP in the BA case shows that the DL models integrate information from the whole brain (**Fig. 5**). However, we see also that neurostructural properties specific to individual participants are detected, specifically in the cortical surface areas, around ventricles, the corpus callosum, at the surface of the brain stem, and cerebellum, and distinct smaller regions in WM areas of the cortex. Ventricles are known to increase in size with age due to regional or global brain shrinkage *(Earnest et al. 1979)*. Also, cortical surface *(LeMay 1984; Kochunov et al. 2005; Jin et al. 2018)*, the corpus callosum *(Doraiswamy et al. 1991)*, cerebellum and basal ganglia *(Raz et al. 2005; Raz et al. 2010)* among others are subject to alterations. While *Raz et al. (2005,* 2010) found no age-related volume changes in, e.g., primary visual cortices and putamen, our model showed that both areas were relevant for the BA estimation across the full life-span, and age-independent rate-of-aging (DBA) in the older cohort (age ≥ 50 years). This may have several reasons: in contrast to linear feature selective models (such as those using regional volume in *Raz et al. 2005; Raz et al. 2010*), our DL-architectures are trained end-to-end, and thus can incorporate information from diverse neural features, including volume, but also region-specific sizes and shapes, tissue structures etc. Within our model those features can be non-linearly related and weighted, and lastly, our multi-modal MLENS leverage this capacity by incorporating complementary image-contrasts.

Similarly, in contrastive relevance maps, we found that heightened DBA values for subjects with type 2 diabetes displayed regions that corresponds to findings of recent animal models *(Muramatsu et al. 2018)* and known diabetes-associated degenerations in the sensorimotor areas in humans *(Ferris et al. 2020)*. Moreover, our results support previous findings in diffusion imaging studies of changes in fiber bundles of the cingulum *(Hoogenboom et al. 2014; Cui et al. 2020)* and neighboring corpus callosum *(Yu et al. 2019)*. That these findings appeared only significant in T1-weighted images, and not, as expected in FLAIR, might be due to the small sample size in the hold-out subset in combination with the less specific contrast of FLAIR in the absence of lesions. However, we found a strong spatial correspondence between the t-maps of both modalities.

We conclude that normal and pathologically driven aging is not exclusively represented in selective features (e.g., in the decline of regional volume) but also in diverse neurostructural properties accentuated by different MRI sequences, throughout the whole brain. More specifically, our analysis pipeline revealed that an individual’s structural MRI carries not only global, macrostructural hints towards its age trajectory, but also reliable information on age-related, subtle grey and white matter changes, including WM lesions that occur all over the brain. While the limited image resolution does not offer explanations at the cellular level, those ubiquitous, rather subtle changes stem most likely from micro-changes, including oxidative stress, DNA damage, cell death and inflammation, in neuronal, vascular and glial compartments of the brain *(Cole and Franke 2017; Pluvinage and Wyss-Coray 2020)* that eventually alter the magnetic properties and thus image contrasts of the respective sequences. We can further infer that all brain regions and different neural properties that are highlighted with the different MRI sequences are predictive w.r.t. age, i.e., the aging process emerges in all these modalities. This calls for a multi-modal approach towards brain-aging rather than restricting this foundational phenomenon to selective neural variables such as grey matter volume, and acknowledges the capability of common structural MRI to reveal not only gross anatomical changes but also subtle microstructural changes with advancing age.

### 4.3. The benefit of multi-level ensemble models

Both types of MLENS performed close to the state-of-the-art in the domain of BA prediction. Note that small performance differences might stem from our smaller dataset with a large age-range in comparison to studies that used, e.g., *UK Biobank* data (n > 14,000 MRIs, age range 44-81 years; e.g., the state-of-the-art model of *Peng et al., 2021*, for brain-age predictions achieved a MAE of 2.14 years. In **Fig.2** of their publication, we can estimate a MAE of 3.1 years for a similar amount of training data, as we used in our study. This performance is almost *on par* with our model, MLENS type 2: MAE = 3.37. The difference is most likely explained by the smaller age-range in the *UK Biobank*). With our MLENS we demonstrated that i) ensembles are performing better than their base models, and ii) MLENS integrating diverse input features, here MRI sequences and brain regions, perform even better than ensembles that are only trained on one of these features.

On a methodological side, this shows that due to the feature selective training the model is prone to specialize on properties inherent to the respective feature (e.g., a brain region). Splitting the brain in sub-regions and feeding them to different models seems to push the respective models (here MLENS type ii) to specialize on the characteristics of each brain region rather than learning filters that are generally usable across the whole brain, however, this needs to be tested systematically.

The variability of predictions between different DL models (here defined as the uncertainty between base models, which was higher for age groups with less MRI data) with an identical architecture and training on the same data, underlines the importance of the aggregation over a set of models (i.e., an ensemble) to reduce both the variance and biases of single networks. In summary, MLENS can not only compensate for the stochasticity of single DL models, but also provide estimates of model certainty and insights on the relationship of input features and prediction.

### 4.4. Brain-age predictions and their association with other biomarkers

To investigate biological determinants of BA, we showed in an additional exploratory analysis that DBA was associated with cardiovascular risk factors such as BMI, waist-to-hip-ratio and type 2 diabetes. Notably, we found that many of these associations depend on the age of participants. For instance, despite the smaller sample size in our younger (healthy) cohort, we discovered a high correlation between BMI and the estimated BA (age < 40 years), which was also reported in *(Kolenic et al. 2018)* for younger participants with first-episode psychosis (18-35 years). Also in mid-aged participants (40-60 years) we saw a significant correlation, for whom previous studies found higher BMI to be associated with cortical thinning *(Shaw et al. 2018)*. Similar to previous findings *(Kharabian Masouleh et al. 2016)*, also in the older cohort (60-80 years), a positive relationship appeared. Overall, with age the association between BA and BMI becomes weaker. Also, we found the positive correlation between DBA and type 2 diabetes, which was reported in *Franke et al. (2013)*, and the corresponding relevance map analysis showed overlapping evidence w.r.t. GM changes as discussed above. Blood glucose levels (here HbA1c) showed relative consistent association across the cohort. With the estimates of MLENS type i this association could also be seen in the 20-35 years old, a result that corresponds to recent findings showing a negative relationship HbA1c and WM integrity in young, non-diabetic (i.e., healthy) adults (mean age 28.8 years, HbA1c < 5.7%; *Repple et al. 2021*), motivating further investigations. Overall, we found similar relationships of DBA and various clinical markers as summarized in *(Franke and Gaser 2019)*, but not regarding ApoE-4 (cf. *Raz et al. 2010*). The found association between DBA and gender should be taken with caution, since demographic factors might have influenced the cohort composition in different age groups. Also, the gender difference is typically most pronounced in younger ages *(Gur et al. 2002)*, while with menopause it appears to become smaller, brain-region specific (e.g., *Ritchie et al. 2018; Raz et al. 2010*) or is even absent *(Jäncke et al. 2015)*. While we found a consistent, slightly negative trend (age > 25 years) between DBA and cognitive performance, the correlation was not significant for most age strata; however, this association has been reported to be more pronounced in patients with AD or mild cognitive impairments *(Gaser et al. 2013; Liem et al. 2017)*. Note that we excluded participants with AD and other neurodegenerative diseases from this study, in which the relationship of DBA to cognitive performance, but also to associated biomarkers such as ApoE14 (see above) might be more pronounced. A very robust positive correlation, nearly across the full age range was found between the WM lesion-load and DBA. The typical accumulation of WM lesions with higher age as well as their pathological consequences are widely known *(Beck et al. 2021; Dinsdale et al. 2021)*, and consequently and conversely, validates the BA models, while in parallel, this highlights the possibility that *typical* and *pathological* aging share similar fundamental mechanisms.

Clearly, these results indicate that BA is a reliable imaging marker reflecting biological plausible age-related neural changes. As deviations from the chronological age correlate with known risk factors for brain damage, BA can be considered as a biomarker of the brain health status of a person.

### 4.5. Limitations and future research

Several limitations need to be considered. First, despite the local information we receive with the LRP heatmaps, they do not explain *per se* what the biological mechanisms are that made the respective highlighted area relevant to the model. For instance, when considering relevant voxels around ventricles, we do not know whether a model tracks the size of a ventricle or potentially alterations at the tissue around it, or both. Further developments in interpretation algorithms, such as LRP could allow the detection of interactions between local and global relevance structures and in addition reveal causal relationships beyond correlation. Second, similar to *Levakov et al. (2020)*, we found that aggregating relevance maps compensated for the observed variability between heatmaps of single base models (for a discussion see *Levakov et al., 2020*). However, aggregation techniques can also cause information loss, for instance, not all of the base models within an ensemble might detect all WM lesions in an image. Third, the age distribution in the LIFE MRI dataset is non-uniform, with a majority of participants being 65 to 75 years of age. This introduces a bias in the training dataset. Moreover, many papers on brain-age estimation reported a corresponding prediction bias towards the mean age in the data *(Cole et al. 2017; Beheshti et al. 2019; Smith et al. 2019; Peng et al. 2021)*. This bias we also observed in our stimulation. Although our ensemble architectures compensate for the prediction bias towards the distribution mean, this tendency could not be fully eliminated. Therefore, we used a sliding-window approach in the correlation analysis with other biological markers, which attenuated this bias further. The assessment of the covariate shift (e.g., *Sugiyama et al. 2007*), nonlinear head-models, and over- or undersampling techniques, combined with data augmentation could be further means to tackle this bias. Moreover, it is to be expected that age-related structural changes systematically effect MRI intensity distributions that models can exploit for their predictions; however, our analysis of relevance maps has shown that the models integrate biologically meaningful brain features across all age groups. Fourth, in future research one could run several cluster analyses to find common relevance patterns within, for instance, participants with certain pathologies or between different age groups. These could then be related to interpretable structural properties, such as cortical thickness *(Frangou et al. 2021)*. Finally, the majority of studies cannot afford to scan thousands of participants. To make the presented explanation pipeline more sustainable, one could explore transfer learning techniques to adapt the pre-trained models to smaller datasets and different (target) variables. Since our approach makes it possible to combine information from different modalities and single out regions which show alterations in these modalities, one might also extend it to incorporating further imaging measures, e.g., diffusion imaging or resting-state studies in fMRI or EEG.

## 5. Conclusion

While certain brain areas shrink in volume more dramatically with older age than others, aging processes emerge in the whole brain. Their progress and pace can now be accurately captured and interpreted by DL ensembles from various brain regions and structural MRI modalities (T1, FLAIR, SWI), proposing that higher age and the presence of cardiovascular risk factors contributes to regionally pronounced yet ubiquitous changes in the brain. Employing the LRP interpretation algorithm, estimates of brain-aging can thus be related back to established, gross but also subtle, most likely microstructural biological markers of the aging process. This bias-free computational approach yields insights into the global nature of brain aging as well as pathomechanisms. Finally, due to its generalizability, this approach can be broadly applied across clinical neuroscience, galvanizing the generation of data-driven hypotheses and boosting its applications in personalized medicine *(Esteva et al. 2021; Stenzinger et al. 2021; Binder et al. 2021)*.

## Acknowledgements

This work is supported by the European Union, European Regional Development Fund as part of the LIFE-LIFT project, and the Free State of Saxony within the framework of the excellence initiative, and LIFE-Leipzig Research Center for Civilization Diseases, University of Leipzig (project numbers 713-241202, 14505/2470), and by the German Research Foundation (project numbers 209933838 CRC1052 Obesity mechanisms A1 and WI 3342/3-1). Further support was provided by the German Ministry for Education and Research (BMBF) through Berlin Institute for the Foundations of Learning and Data (BIFOLD; refs. 01IS18025A and 01IS18037A), MALT III (ref. 01IS17058), Patho234 (ref. 031L0207D) and Transparent Medical Expert Companion (TraMeExCo, ref. 01IS18056A), European Union’s Horizon 2020 research and innovation programme through Intelligent Total Body Scanner for Early Detection of Melanoma (iToBoS, grant agreement No 965221), as well as the Grants 01GQ1115 and 01GQ0850; and by Deutsche Forschungsgemeinschaft (DFG) under Grant Math+, EXC 2046/1, Project ID 390685689; by the Institute of Information & Communications Technology Planning & Evaluation (IITP) grant funded by the Korea Government (No. 2019-0-00079, Artificial Intelligence Graduate School Program, Korea University).

## Author Contributions

S.M.H., K.R.M., A.V., W.S., and A.V.W. designed and discussed the project.

Responsible for data acquisition M.L., A.V., and A.V.W.

Data analysis was done by S.M.H., O.G. and F.B.

Writing and editing were done by S.M.H., F.B., S.L., K.R.M., A.V., and A.V.W.

S.M.H. made the figures and tables.

