## Appendices for "Towards the Interpretability of Deep Learning Models for Multi-modal Neuroimaging: Finding Structural Changes of the Ageing Brain"

### Appendences

#### Appendix A. Supplementary Methods

##### MRI processing stages

***Raw stage*** raw MRI DICOMs were saved in the NifTi-1 format in their native space.

***Freesurfer volume stage*** First, T1-MRIs were subject to the recon-all preprocessing pipeline of FreeSurfer 5.3.0 *(Fischl 2012)*. This includes a brain extraction step, intensity normalization procedures, linear registration to the FreeSurfer standard space, and rescaling between 0-255. Then, the intermediate processing stage of the T1-image ('brain.finalsurf.mgz') was used to linear register (Rigid, linear interpolation; ANTs 2.2; *Avants et al. 2011*) also images of the other two sequences (FLAIR, SWI) to the space of the T1-weighted images. Before rescaling the images of these two MR sequences to the same range (0-255), high intensity *outliers* were clipped to 383, which guaranteed that their intensity distributions do not show skewed biases based on high intensity noise nor high intensity values which potentially correlate with age, e.g., in white matter lesions.

***MNI stage*** To bring images also to a common space across participants, all available sequences were non-linearly warped to the MNI152 space *(Fonov et al. 2011)* with 2 mm isotropic resolution (ANTs 2.2).

***Freesurfer surface stage*** This stage is a targeted output of FreeSurfer's *recon-all* pipeline, mapping the brain in volume-space to surface-space by creating a 3D-mesh around its folds.

The corresponding computed mapping files were later used to first convert and then explore the relevance maps of our interpretation algorithm (LRP) in the individual surface space. Hence, this image stage was only used for visualization and analysis after the training of the prediction models.

##### Brain atlases

For the model training on distinct regions of the brain (multi-layer ensemble, MLENS type ii), as well as for the structural mapping of LRP relevance distributions, we used a combination of three atlases that cover nearly the entire brain as defined by the MNI152 template: the *Harvard-Oxford* i) cortical and ii) subcortical structural atlases, and iii) the cerebellar atlas *(Diedrichsen et al. 2009)*, all distributed via *FSL 5.0.8*. While there is a minimal overlap between the cerebellum to the other two atlases, we removed the left and right cerebral cortex labels from the subcortical atlas, due to their informational redundancy with respect to the cortical atlas. For details on the Juelich histological atlas, see: *https://fsl.fmrib.ox.ac.uk/fsl/fslwiki/Atlases/Juelich.*

#### Appendix B. Additional simulation studies

We ran two further simulations to test how the model captures i) shape and ii) gradual intensity shifts as functions of age, respectively. For the first simulations, age was introduced as linear changes in height and width of a torus. Young tori are taller and thinner, while old tori become flatter and wider. In contrast to the main simulation, lesions and atrophies were not added. For the second additional simulation, the shape of tori was kept equal across the full age range, adding a degree of randomness; also, no lesions nor atrophies were added. Instead, age was represented as a linear gradual intensity shift including random noise in the upper quadrant of a torus. Young tori have low intensity values, while older tori have higher values in that quadrant (maximum at 1; for details see the open repository XDLreg;

*https://github.com/SHEscher/XDLreg*). All other hyperparameters, such as model architecture, training epochs etc., where the same as in the main simulation.

In both simulations, the respective model could predict age with a high accuracy in the unseen test set (simulation i: MAE = 1.69; simulation ii: MAE = 2.36). For the shape simulation, the LRP analysis revealed that the model particularly *looks* at the boarder of a torus for its age estimation (**Fig. A1a**). In the simulation of the age-related local intensity shifts, the LRP analysis revealed that the model primarily focuses on the respective upper quadrant (**Fig. A1b**).

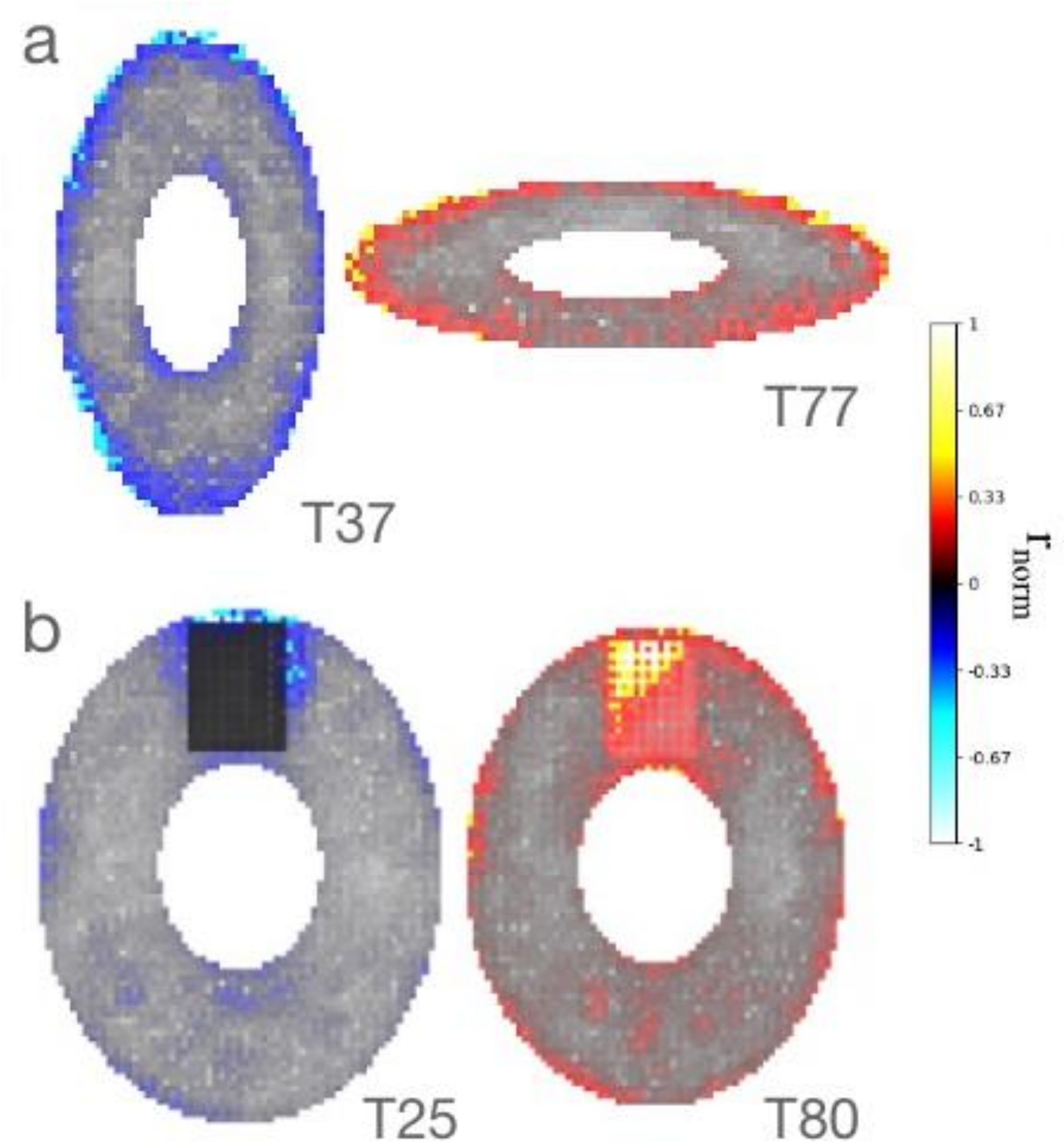


***Fig. A1 (a) Shape simulation*** *The age of tori is purely a function of their shape. No atrophies nor lesions are added.* Left*: Torus of age 37, predicted 33.7.* Right*: Torus of age 77, predicted 74.9. LRP reveals that the model mainly looks at the border of tori, potentially estimating height and width, which both reflect age in this additional simulation.* ***(b) Local intensity simulation*** *Here the age of a torus is a function of the intensity values in the upper quadrant of its body. The shape of tori across the full age range is equal, adding some randomness. No atrophies nor lesions are added.* Left*: Torus of age 25, predicted 21.7.* Right*: Torus of age 80, predicted*

*74.9. LRP reveals that the model mainly looks at upper quadrant of a torus, which reflects age in this simulation. However, also some random locations at the border and in the body of the tori are picked up. Note that the darker a location is (here upper quadrant of left torus) the less strong relevance values are visible. This is because voxel intensities are multiplied with relevance values for visualization. Consequently, relevance values in the quadrant of the young tori appear more dark than blue.*

### Appendix C. Model prediction performance

| **Ensembles** | **Head model** | **Base models** | | | |
|---|---|---|---|---|---|
| | | **$mean_{MAE}$±SD** | **$min_{MAE}$** | **$max_{MAE}$** | **$N_{bm, MLENS}$** |
| **Multi-level ensemble (type i)** | **3.88** | - | - | - | **30** |
| T1 sub-ensemble | 4.31 | 5.15±0.94 | 4.42 | 13.89 | 10 |
| FLAIR sub-ensemble | 4.13 | 5.12±1.53 | 3.99 | 12.93 | 10 |
| SWI sub-ensemble | 5.83 | 6.55±0.87 | 5.15 | 13.44 | 10 |
| **Multi-level ensemble (type ii)** | **3.69** | - | - | - | **45** |
| Cortical-T1 sub-ensemble | 5.10 | 6.89±2.61 | 4.51 | 14.77 | 5 |
| Cortical-FLAIR sub-ensemble | 4.34 | 6.36±2.89 | 4.34 | 13.52 | 5 |
| Cortical-SWI sub-ensemble | 6.11 | 7.66±1.78 | 5.44 | 14.77 | 5 |
| Sub-Cortical-T1 sub-ensemble | 9.88 | 12.22±2.06 | 6.42 | 14.78 | 5 |
| Sub-Cortical-FLAIR sub-ensemble | 5.34 | 8.91±3.97 | 4.48 | 12.84 | 5 |
| Sub-Cortical-SWI sub-ensemble | 6.33 | 9.01±2.54 | 5.67 | 14.77 | 5 |
| Cerebellum-T1 sub-ensemble | 6.28 | 9.45±3.22 | 5.72 | 14.53 | 5 |
| Cerebellum-FLAIR sub-ensemble | 4.89 | 6.52±2.51 | 4.98 | 14.53 | 5 |
| Cerebellum-SWI sub-ensemble | 7.58 | 8.80±1.18 | 7.03 | 14.77 | 5 |

***Table A2 Prediction performances of both multi-level ensembles (type i, ii) with ReLU activation functions** and their respective sub-ensembles and 3D-CNN base models (bm), measured in mean absolute error (MAE). Note, MAEs > 13 indicate that a specific model did not learn from the data, i.e., it only output the approximate population mean, which was set at the bias in the output layer.*

#### Appendix D. Data distribution of LIFE

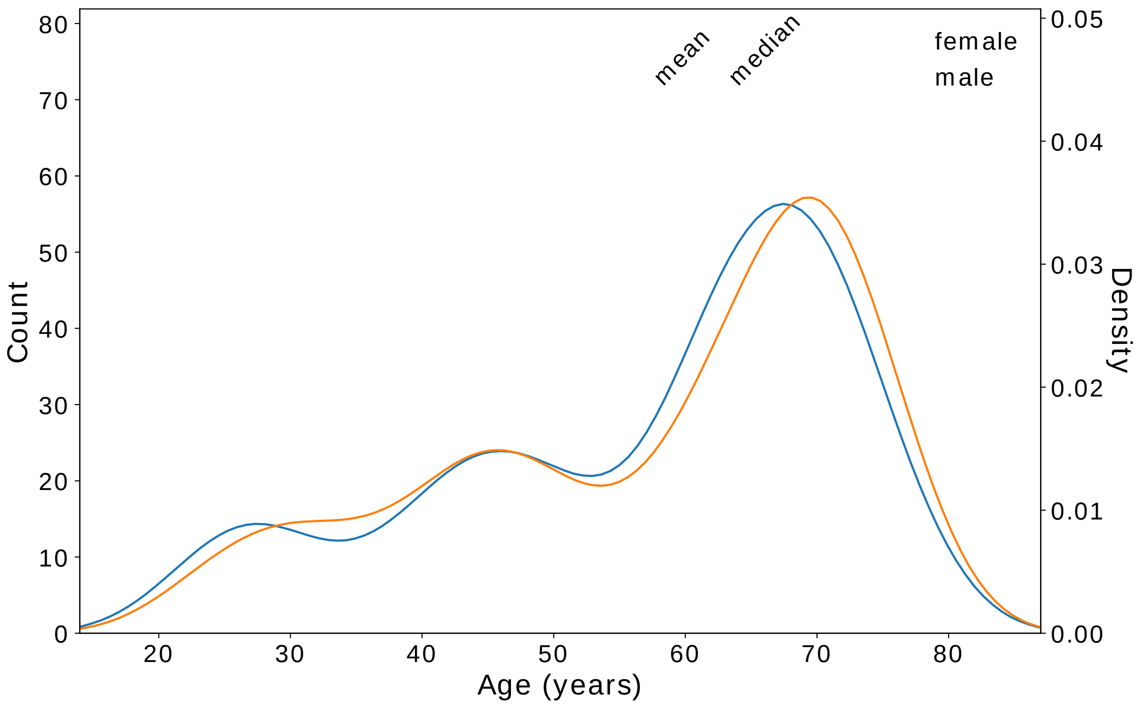


***Fig. A3 Age distribution in LIFE MRI dataset*** *after exclusion (n=2016, mean age = 57.32, median age = 63.0).*

#### Appendix E. Relevance per brain component

For each participant, relevance values from the aggregated relevance map of MLENS type i were summed in three brain components (grey matter, GM; white matter, WM, cortical spinal fluid, CSF), respectively, and plotted over the corresponding ages of the participants (**Fig. A4**). To retrieve the relevance per component, relevance maps were warped to MNI152 space (2mm resolution) and then masked with the respective component using *nilearn 0.9.0* (specifically the functions: *load_mni152_gm_mask* for GM, *load_mni152_wm_mask* for WM, and *fetch_icbm152_2009* for CSF). For nearly all components, but CSF in the SWI sub-ensemble, there was a significant correlation between age and the sum-score (see **Table A5**). Note that, here, the sum of relevance only approximates the overall prediction, since i) it is split in three components, and ii) they are taken from the aggregated relevance map of single participants (i.e., the average relevance map between base models of a sub-ensemble; see **Section 2.5.2.** in the main manuscript).

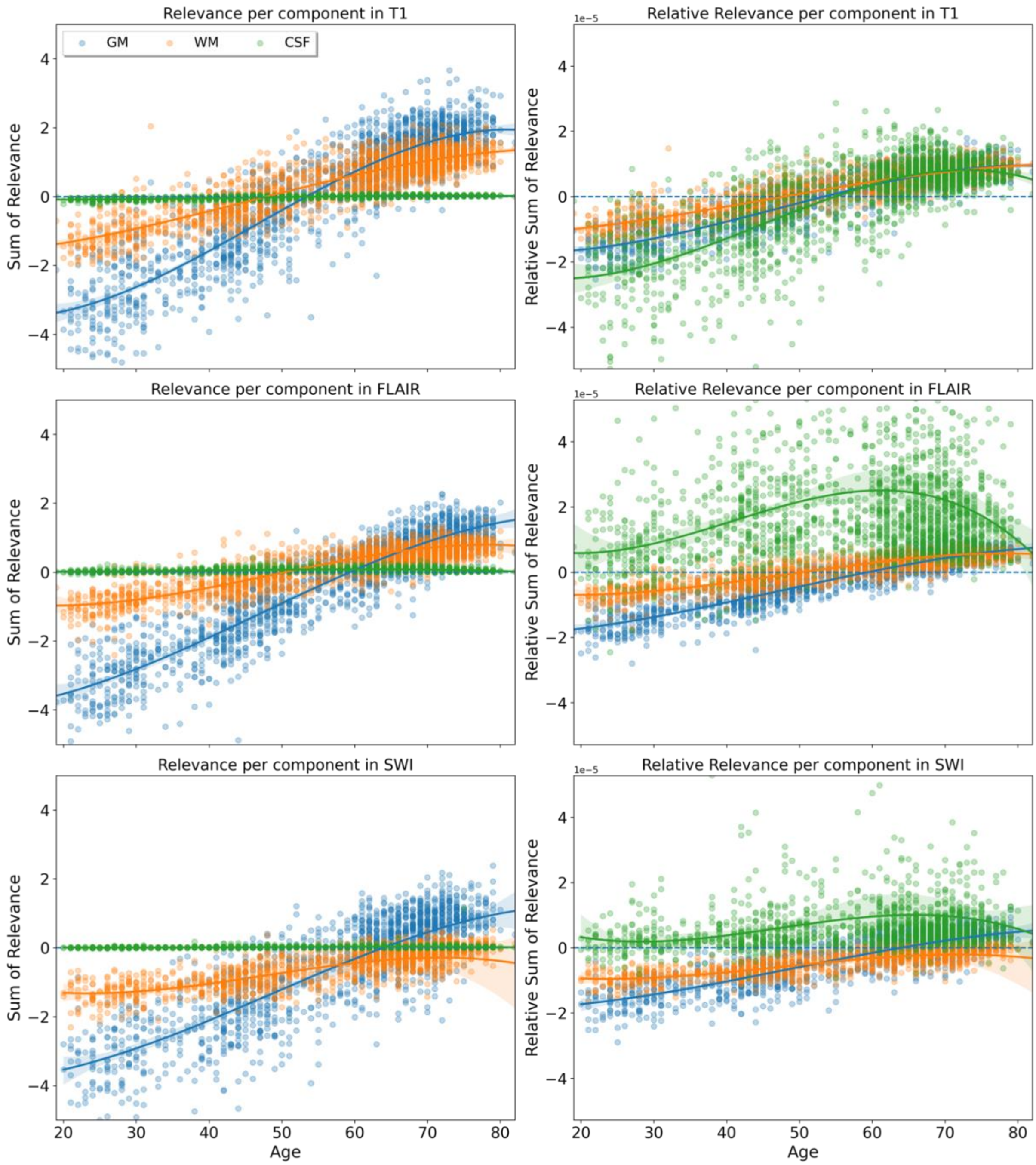


***Fig. A4 Summed relevance per brain component*** Left column*: sum of relevance values over components (grey matter: GM, white matter: WM, cortical spinal fluid: CSF).* Right column*: Relative sum of relevance over components, normalized by the respective size (in number of voxels) of each component.*

| | GM | WM | CSF |
|---|---|---|---|
| **T1** | *R* = 0.91, *** | *R* = 0.88, *** | *R* = 0.76, *** |
| **FLAIR** | *R* = 0.50, *** | *R* = 0.27, *** | *R* = 0.05, * |
| **SWI** | *R* = 0.49, *** | *R* = 0.16, *** | *R* = 0.05, *ns* |

***Table A5 Correlation table of summed relevance per brain component and age*** *Pearson's correlation (R) between age and sum of relevance values in the three major brain components grey matter (GM), white matter (WM) and cortical spinal fluid (CSF) in all three sub-ensembles of MLENS type i that were trained on different MRI sequences (T1, FLAIR, SWI). ns: p>0.05; *: p<0.05; **: p<0.01; ***: p<0.001.*

### Appendix F. Relevance difference between MRI modalities

To explore how the models extract age information differently between MRI modalities, we took the aggregated relevance maps from the sub-ensembles of MLENS type i, that were trained with MRIs in one of the three modalities (T1, FLAIR, SWI), respectively. For each participant, we subtracted the relevance maps in a pair-wise fashion from each other (T1-FLAIR, T1-SWI, FLAIR-SWI). We then run a *1-sample t-test* on these three difference maps over the full age-range using *FSL 5.0.8* (*randomise* function; 5000 permutations; threshold-free cluster enhancement). As expected, we find that the model predominantly attributes higher relevance to cortical grey matter areas, including the occipital lobe in T1-weighted images than in FLAIR and SWI. In contrast, white-matter in sub-cortical but also in fronto-cortical areas showed higher relevance in FLAIR in comparison to T1. In SWI, the model attributed higher relevance in ventral parts of the brain including the brain stem and ventricles towards the cerebellum, but also bilateral regions of the central opercular cortex in contrast to T1-weighted images (**Fig. A6**).

Note, here, higher relevance is relative: For individual regions there can be age-related biases within the sub-ensembles. For instance, if the T1 sub-ensemble looks in a specific region exclusively for information towards a younger age, that is, attributing only negative relevance values to this region if applicable, while the FLAIR sub-ensemble ignores the region for all of its predictions (zero relevance), this would lead to a negative difference score in that region (T1-FLAIR). In turn, the corresponding difference map could be misinterpreted by falsely assuming that the specific region is more informative in FLAIR (towards a higher age), due to the flipped sign in the difference maps in that particular region.

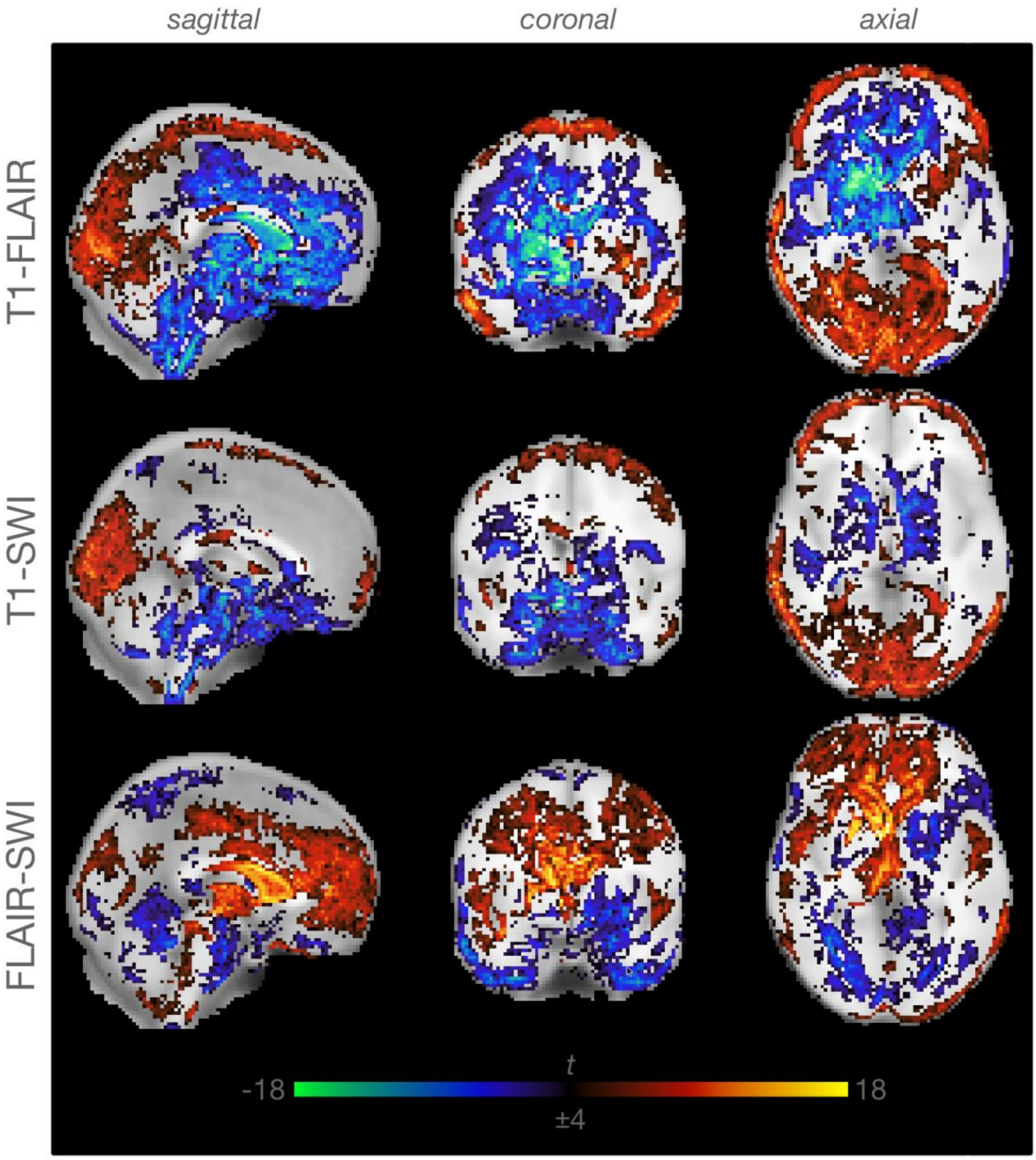


***Fig. A6 Difference relevance maps of three MRI sequences T1, FLAIR, SWI*** *After subtracting relevance maps per participant in different MRI sequences from each other, 1-sample t-tests were computed to find areas which showed significant differences between the*

*modalities.* Rows*: Three difference maps (T1-FLAIR, n=402; T1-SWI, n=314; FLAIR-SWI, n=314), which were cut (x=0, y=0, z=0) in three orientations (*columns*).* Red-yellow *colors indicate where relevance values are higher in the first modality in contrast to the second modality, and vice versa for* blue-green *colors (e.g., i. T1 and ii. FLAIR, in the first row T1-FLAIR). T-maps were clipped at (-18, -4) and (4, 18).*

### Appendix G. Contrastive relevance maps

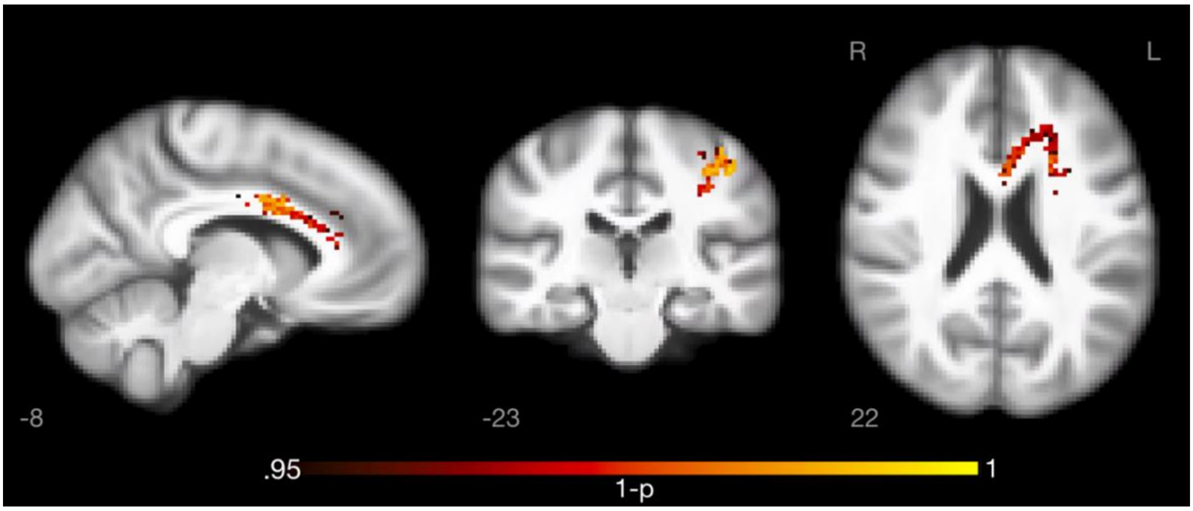


***Fig. A7 Contrastive relevance maps of diabetics vs. control****. For T1-sub-ensemble in MLENS type I, subjects with type 2 diabetes mellitus (n=29) were contrasted to controls (n=217; TFCE, FWE- corrected p ≤ 0.05) in older subjects (50-75 years).*

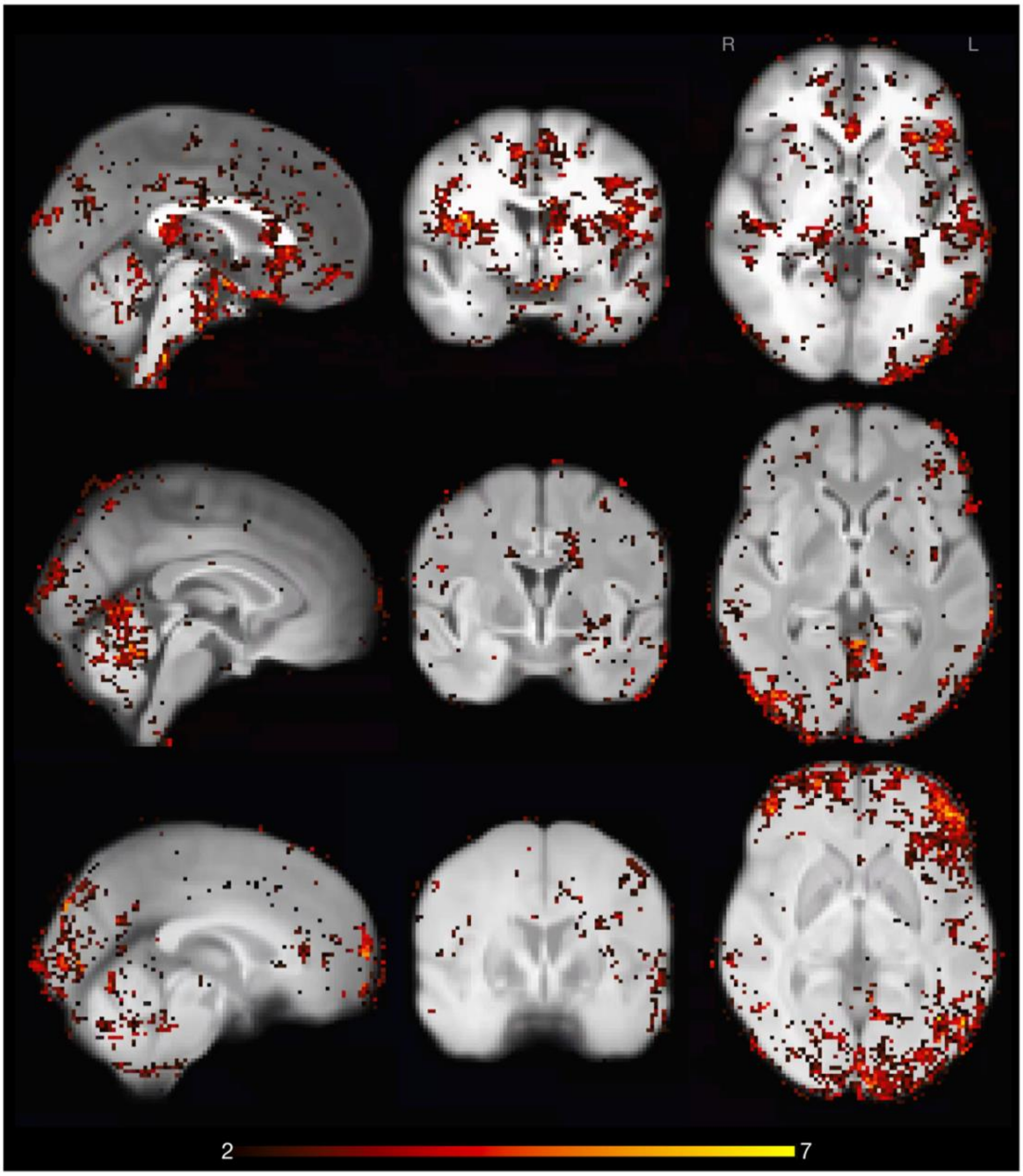


***Fig. A8 The role of diverging brain age (DBA) on relevance attribution*** *T-maps (2, 7) of the GLM analysis on the modulation of relevance maps as function of DBA, corrected for age in the older sub-cohort (age ≥ 50).*

### Appendix H. Age-bias corrected diverging brain-age

In spite of the architectures of multi-level ensembles, we still found an age-bias in diverging brain-age (DBA; **Fig. 2**), which also other studies have reported (e.g., *Cole et al., 2017; Peng et al., 2021; Smith et al., 2019*). This bias might affect the correlation analysis between DBA and other bio- and lifestyle-related markers (**Fig. 7**) despite the applied sliding-window approach, which aimed to minimize this effect. To attenuate the role of the age-bias in DBA further, we fitted a linear model between age (independent variable) and DBA (dependent variable), and then subtracted the newly estimated DBA* of the linear model from the original predictions, similar to *Beheshti et al. (2019)*. In contrast to these authors, we used a polynomial fit, i.e., with a degree > 1. We found that the polynomial degree of 4 showed the highest correlation between age and DBA across the full sample (MLENS type i, $R^2$=0.14; MLENS type ii, $R^2$=0.14); that is, it was optimal to minimize the respective mean absolute errors (MAEs) of the two multi-level ensembles (MLENS type i, MAE=3.56, cf., uncorrected MAE=3.86; MLENS type ii, MAE=3.05, uncorrected MAE=3.37). Note, we explicitly overfitted the linear *correction* model (i.e., we did not fit its coefficients on separated data) in order to maximally reduce the effect of age in the DBA correlation analysis.

For the correlation analysis between the bias-corrected DBA and other variables, we found that the overall trends remained the same (**Fig. A9**, cf. **Fig. 7**). For some variables the number of significant correlations (per window) with the bias-corrected DBA remained in a narrower age-range, for instance, in BMI between 35-45 years, which showed before significant correlations up to 70 years. In contrast, for WM lesions the correlations in the oldest population were even more pronounced. This was to be expected, since the initial age-bias in DBA resulted in older people being estimated on average to be younger than they actually are (**Fig. 2**), while there is an expected increase of WM lesions *(Beck et al. 2021)*.

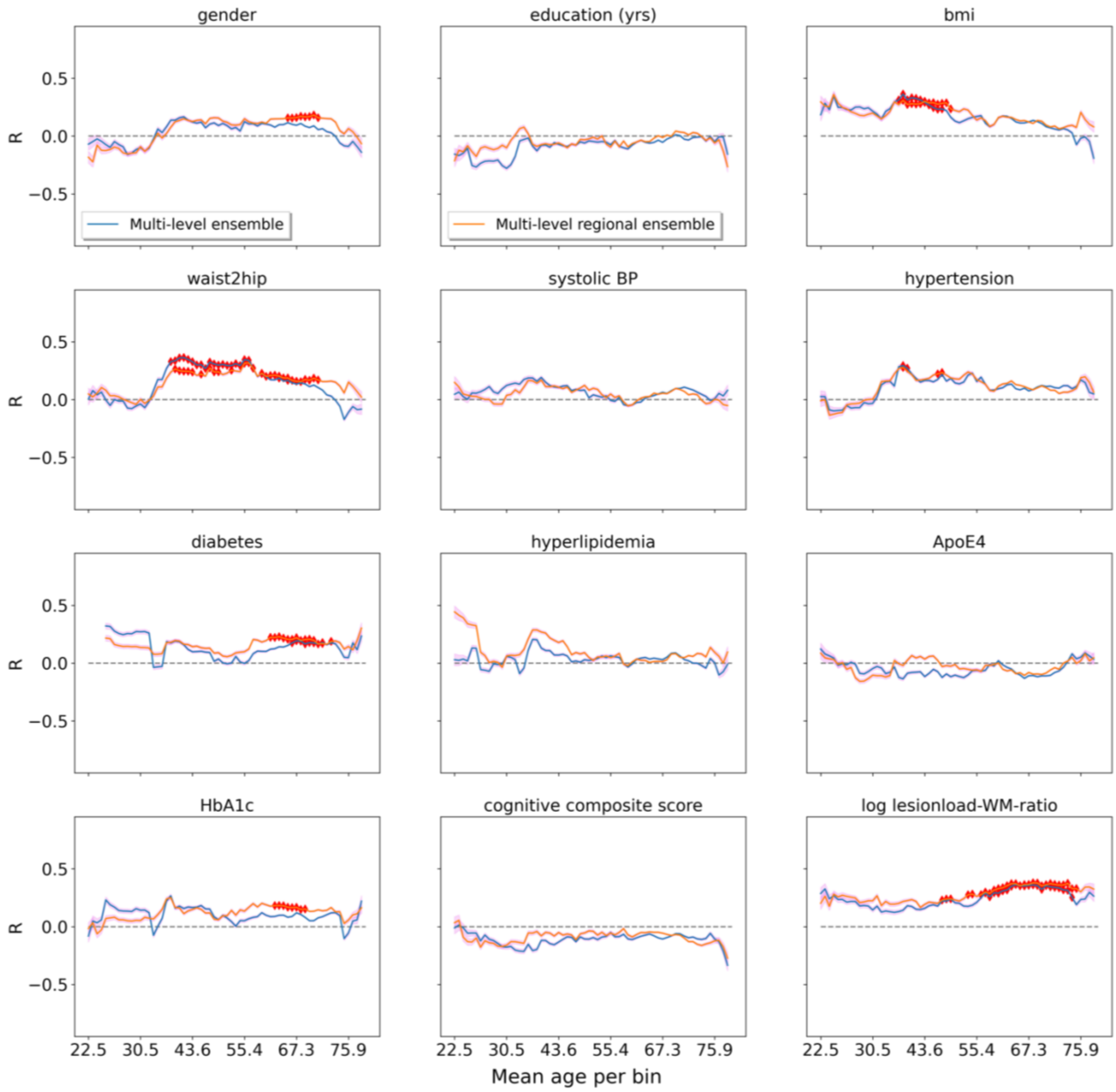


***Fig. A9 Relationship of diverging brain-age (DBA) to biomarkers and lifestyle factors after correcting DBA for its age-bias*** *Similar to* ***Fig.7****, with the difference that DBA (i.e., prediction error) is corrected for the observed age-bias* (regression towards the mean, ***Fig. 2****). After the age-correction, the overall trend in all variables remains similar. For some variables the number of significant correlations in age-windows is reduced to a narrower age-range, as in bmi, whereas in other variables there is a shift towards an older age, as in log lesionload-WM-ratio.*

### Appendix I. References

Avants BB, Tustison NJ, Song G, Cook PA, Klein A, Gee JC. 2011. A reproducible evaluation of ANTs similarity metric performance in brain image registration. NeuroImage. 54(3):2033–2044. doi:10.1016/j.neuroimage.2010.09.025.

Beck D, de Lange A-MG, Maximov II, Richard G, Andreassen OA, Nordvik JE, Westlye LT. 2021. White matter microstructure across the adult lifespan: A mixed longitudinal and cross-sectional study using advanced diffusion models and brain-age prediction. NeuroImage. 224:117441. doi:10.1016/j.neuroimage.2020.117441.

Beheshti I, Nugent S, Potvin O, Duchesne S. 2019. Bias-adjustment in neuroimaging-based brain age frameworks: A robust scheme. NeuroImage: Clinical. 24:102063. doi:10.1016/j.nicl.2019.102063.

Cole JH, Poudel RPK, Tsagkrasoulis D, Caan MWA, Steves C, Spector TD, Montana G. 2017. Predicting brain age with deep learning from raw imaging data results in a reliable and heritable biomarker. NeuroImage. 163:115–124. doi:10.1016/j.neuroimage.2017.07.059.

Diedrichsen J, Balsters JH, Flavell J, Cussans E, Ramnani N. 2009. A probabilistic MR atlas of the human cerebellum. NeuroImage. 46(1):39–46. doi:10.1016/j.neuroimage.2009.01.045.

Fischl B. 2012. FreeSurfer. NeuroImage. 62(2):774–781. doi:10.1016/j.neuroimage.2012.01.021.

Fonov V, Evans AC, Botteron K, Almli CR, McKinstry RC, Collins DL. 2011. Unbiased average age-appropriate atlases for pediatric studies. NeuroImage. 54(1):313–327. doi:10.1016/j.neuroimage.2010.07.033.

Peng H, Gong W, Beckmann CF, Vedaldi A, Smith SM. 2021. Accurate brain age prediction with lightweight deep neural networks. Medical Image Analysis. 68:101871. doi:10.1016/j.media.2020.101871.

Smith SM, Vidaurre D, Alfaro-Almagro F, Nichols TE, Miller KL. 2019. Estimation of brain age delta from brain imaging. NeuroImage. 200:528–539. doi:https://doi.org/10.1016/j.neuroimage.2019.06.017.
